# Lipid Nanoparticle Delivery of Mesenchymal Stromal Cell-Derived microRNA 187-3p as a First-in-Class Therapy for Myocardial Dysfunction in Sepsis

**DOI:** 10.1101/2025.09.09.674991

**Authors:** Amin M. Ektesabi, Maria Carolina Barbosa-Silva, James N. Tsoporis, Samantha McWhirter, Chirag M. Vaswani, Yanis Mouloud, Logan W.C. Zettle, Abdalla Ahmed, Paul Delgado-Olguin, Sabrina Setembre Batah, Jean-Francois Desjardins, Sahil Gupta, Greaton W. Tan, Jacqueline E. Pavelick, Yanbo Wang, Alexandre T. Fabro, Phyllis Billia, Nikolaos Kavantzas, John C. Marshall, Kim A Connelly, Shirley H.J. Mei, Lauralyn McIntyre, Duncan J. Stewart, Howard Leong-Poi, Tatiana Maron-Gutierrez, Bernd Giebel, Gilbert C. Walker, Claudia C dos Santos

**Author notes:** Correspondence and reprint requests to: C. C. dos Santos, MSc, MD., Clinician-Scientist, Associate Professor of Medicine, Interdepartmental Division of Critical Care, St. Michael’s Hospital/University of Toronto, 30 Bond Street, Room 4-008, Toronto, ON, M5B 1WB, CA. Both authors contributed equally to this work (Amin M. Ektesabi, Maria Carolina Barbosa-Silva). **Acknowledgements:** We acknowledge our membership in the Canadian Critical Care Translational Biology Group (CCCTBG, CCDS, JM), Sepsis Canada (JM and CCDS), the Nanomedicine Innovation Network (GW), and the International Society for Gene and Cell Therapy (CCDS, DJS, SM, BG). We further acknowledge the CFI John R. Evans Leaders Fund (CFI-JELF), St Michael’s Hospital Foundation and biobank manager Dr. Valeria DiGiovanni for support of the creation and management of the Precision Medicine for Critical Care (PREDICT)-biobank at Unity Health Toronto. We especially thank the support of the Keenan Center for Biomedical Research vivarium (veterinary doctor, managers and animal technicians), the technician and managers at the core facility (Drs. Caterina Di Ciano-Oliveira, Monika Lodyga, Pamela Plant) as well as the research coordinators (Marlene Santos and Gyan Sandhu) and laboratory technicians (Eryn Churcher and Shelah Izhar) at the Critical Care Research Unit and dos Santos Lab for their ongoing support of our translational work. **Sources of Funding:** Our work has been funded by grants from the Canadian Institutes of Health Research (CIHR, CCDS, DJS, JCM, GW, PDO), the Nanomedicine Innovation Network (GW, CCDS), CCDS held the Robert and Dorothy Pitts Research Chair in Acute and Emergency Medicine and now holds the Tier 1 Canada Research Chair for accelerated Molecular Critical Care. MITACs internship grants (NorthMiRs, CMV, AME, SW, LZ), PhD award scholarship from Brazilian funding agency CAPES (Coordenação de Aperfeiçoamento de Pessoal de Nível Superior) and ISN (International Society for Neurochemistry) and Post-Doctoral fellowship awards to MCBS (Keenan Research Center for Biomedical Research), CMV (CIHR), University of Toronto Emerging and Pandemic Infections Consortium Convergence Postdoctoral Fellowships (AME), and Ted Rogers Centre for Heart Research Education Fund’s PDF and PhD (AME, AA), Emerging and Pandemic Infections Consortium (EPIC) PhD award (GT), Fundação de Amparo à Pesquisa do Estado de São Paulo (FAPESB) PhD award from Brazil (SSB), University of Toronto Queen Elizabeth Award MSc (JP), and Angels Den (St. Michael’s Hospital Foundation). **Disclosures:** CCDS, GW, SW, LZ, AME, JT and CMV are co-founders and shareholders of NorthMiRs that have licensed patents developed for this study and are actively commercially developing innovative miR-based nanotherapies for sepsis. CMV, AME, SW and LZ are NorthMiRs company interns partially funded by MITACs. BG is the founder and CEO of Exosla Therapeutics. **Author contribution:** LNP-miR187experiments (AME), EV experiments (MCBS and TMG), MSC experiments (JT and JFDJ), LNP/EV codelivery experiments (AME, MCBS), LNP design and production (GW and SM), LNP quality control and in vitro cell toxicity (LZ and JT), ciMSC and EV production (YM and BG), in vitro EV experiments (AME and JT), invasive cardiac monitoring (JFD and JT), human serum and whole blood sample collection and analysis (CCDS, JT, AME, SG, JM), human hearts sample collection and pathology analysis (NK, JT, SSB and AF), RNA sequencing (PD), RNA seq and microRNA seq analysis (AME and CCDS), in-vivo monitoring and welfare assessments, qRT-PCR, ELISAs and ddPCR (AME, MCBS, JT, CMV, GT, JP, YW), echocardiograms and data interpretation (JT, AME, HLP, KC and PB), microRNA scope and analysis (SSB, ATF, JP and CMV), human MSC clinical study, sample collection and analysis (DS, SM, LM), writing of first draft (MCBS, AME and JT), research design and conception (CCDS, BG and GW). All authors read and approved the final draft of the manuscript.

## Abstract

**Background:** Sepsis-induced myocardial dysfunction is a common and critical complication of sepsis. Extracellular vesicles (EVs) from clonally expanded immortalized mesenchymal stromal cells (ciMSCs) contain microRNAs that may be exploited as therapy.

**Methods:** In mouse models of septic cardiomyopathy induced by caecum ligation and puncture, cardiac function was determined by invasive and echocardiographic assessment. Primary cardiomyocytes derived from foetal murine and human adult ventricular tissue, as well as murine hearts were used for mechanistic studies. Studies using post-mortem human hearts or patient plasma, and clinical and echocardiographic measurements were used to establish translational relevance.

**Results:** In preclinical models of sepsis, intravenous administration of either MSCs or ciMSC-EVs, given after the induction of sepsis, prevented a decrease in myocardial ejection fraction, ventricular inflammation, and mortality compared to placebo or platelet-derived control EVs. EV-microRNA sequencing identified enrichment for microRNA-187a-3p (miR-187) in ciMSC-EVs. miR-187 is anti-inflammatory; with interleukin-6 (IL-6) as its major target. Intravenous delivery of lipid nanoparticle (LNP) encapsulated miR-187 improved cardiac function, reduced inflammation, and enhanced survival of septic mice. In cardiomyocytes and in murine hearts, LNP-miR-187 reduces inflammation and expression of myocardial transcription factors linked to fetal gene reactivation in failing septic hearts. In human septic hearts, low circulating miR-187 levels correlate with reduced cardiac function and high sequential organ failure assessment (SOFA) scores.

**Conclusion:** These findings support the development of first-in-class, cell-free, miRNA-based therapy as a novel approach to treat sepsis-induced cardiomyopathy to address a critical gap in sepsis care.

**One Sentence Summary:** miR-based therapy for sepsis

**The Clinical Perspective:**

**A. What is NEW?** Sepsis accounts for 1 in 5 deaths worldwide. Here, we demonstrate that sepsis-induced myocardial dysfunction represents a discrete, targetable sepsis-trait — a distinct biological abnormality characterized by cardiomyocyte inflammation and fetal gene reactivation. This component contributes to the propagation of organ dysfunction and overall mortality and may respond to focused epigenetic-based interventions.
**B. What are the Clinical implications?** Currently, there are no effective treatments to reduce, limit, or reverse the immune dysfunction component of sepsis that contributes to multiorgan failure, such as sepsis-induced cardiomyopathy. We identify miR-187 as a clinically relevant post-transcriptional regulator of cardiac inflammation and cardiomyocyte gene expression. Intravenous delivery of miR-187 encapsulated in a lipid nanoparticle (LNP) represents a fundamentally distinct, effective and pathogen-agnostic approach to correcting sepsis-induced cardiac dysfunction through modulation of cardiomyocyte inflammatory and metabolic pathways.

## Introduction

Sepsis, defined as life-threatening organ dysfunction caused by a dysregulated host response to infection [1], accounts for an estimated 11 million annual deaths globally. The development and persistence of new or pre-existing organ failure, even after pathogen elimination, remains the major cause of mortality and the most clinically important manifestation of sepsis. While antimicrobials and organ-supportive measures have improved survival, specific pharmacological treatments to reduce, limit, or reverse organ dysfunction remain elusive.

Over the past two decades, mesenchymal stromal cells (MSCs) have been investigated as a potential treatment for sepsis. MSCs have unique immunomodulatory, antimicrobial, and bioenergetic resuscitative properties. Much of the therapeutic benefit of MSCs is attributed to their ‘secretome’, composed of an array of bioactive molecules including cytokines, chemokines, growth factors, and extracellular vesicles (EVs). MSC-derived EVs have garnered particular interest for their ability to deliver proteins, lipids, and non-coding RNAs, such as microRNAs (miRNAs), presenting a novel strategy for the development of cell-free therapies for sepsis.

One key target for such therapies is sepsis-induced myocardial depression, a treatable but often overlooked component of sepsis-associated organ dysfunction. Cardiac dysfunction occurs in 30– 60% of patients with sepsis and is characterized by reduced cardiac output and a left ventricular ejection fraction (LVEF) of less than 45% within the first 48–72 hours. When present, sepsis-induced cardiac dysfunction significantly increases the likelihood of death.[2–4] The mechanism by which the heart contributes to mortality goes beyond the detrimental effects of ‘pump-failure’ alone and includes the release of inflammatory mediators from injured cardiomyocytes, such as tumor necrosis factor alpha (TNFα), interleukin 6 (IL-6), and S100 proteins A8/A9/A1, that interact with damage-associated molecular patterns (DAMPs) to propagate immune activation and organ failure as well as myocardial remodeling with reactivation of foetal genes known to also play a role in heart failure.[5, 6] We and others have shown that MSCs and EVs mitigate inflammation and the release of cardiac DAMPs.[7, 8] Various studies indicate miRs contained in MSC-derived EVs (EV-miRs) may confer therapeutic benefit in models of sepsis [9]. We have shown that endogenous miRs are regulated by MSC-administration and transfection of miR mimics or inhibitors, may attenuate bacterial and viral-induced acute lung injury (ALI) and sepsis-induced cardiomyopathy.[10–12] MiR-based therapies target RNA transcripts by altering transcript processing, transcript stability, or translation. The distinct advantages of miR-based drugs include specificity, the ability to target traditionally difficult to hit targets, and the possibility of regulating groups of functionally related RNAs *in situ*.

Here, we hypothesized that delivery of regulatory MSC-derived EV-miRNAs that modulate epigenetic pathways, may modifying the miRNA-mRNA interactome, reduce inflammation and improve cardiac function and survival in sepsis. To test this, we administered MSCs and EVs from clonally expanded and immortalized MSCs (ciMSCs), a source of stable and reproducible EVs, in a murine model of sepsis-induced cardiomyopathy. We sequenced and compared MSC-derived EV-miRs with control platelet-derived EV-miRs to find a candidate therapeutic miR-187a-3p (hereafter referred to as miR-187) significantly enriched in MSC-derived EVs. Amongst differentially expressed microRNAs, miR-187 levels were significantly decreased in circulating and post-mortem septic compared to non-septic hearts. We selected miR-187 mimic for encapsulation in a lipid nanoparticle (LNP) and delivered the LNP-187 formulation intravenously to septic mice, where LNP-miR-187 significantly attenuated myocardial dysfunction, reduced sepsis-associated mortality, and limited fetal gene reactivation in murine hearts. Our data provide preclinical evidence supporting the potential of miR-based nanotherapeutics as a strategy for treating sepsis-induced myocardial dysfunction.

## Results

### Intravenous (IV) MSC administration improved survival and attenuated sepsis-induced myocardial dysfunction

Male C57Bl/6 mice were subjected to cecal ligation and puncture (CLP) or sham surgery and, six hours later, randomized to receive either 2.5×10⁵ human bone marrow–derived MSCs or saline (placebo) via tail vein injection (experiment design is shown in Figure 1A, detailed materials and methods provided in supplement). All mice received antibiotics, fluid resuscitation and pain management. Systemic MSC-delivery improves the early probability of survival (48 hrs) by ∼27% compared to antibiotics alone (saline placebo, Figure 1B). Increased survival was associated with an increase in ejection fraction (18% EF, Figure 1C) and fractional shortening (25% FS, Figure 1D, Figure 1C-E). Invasive hemodynamic measurements performed at 48 hours post-CLP confirmed an 11% improvement in left ventricular (LV) pressure (mean 99 vs. 88 mmHg in placebo - Fig 1F), 24% reduction in LV end-diastolic pressure (LVDP, Figure 1G), a 22% decrease in dp/dt_min_ (Fig 1H), and 27% improvement in dp/dt_max_ (Fig 1I). LV focal myocardial inflammation was similar in all CLP groups as evaluated by quantification of polymorphonuclear (PMN) cell myocardial tissue-area percent-fraction infiltration (Figure 1J).

**Figure 1.**
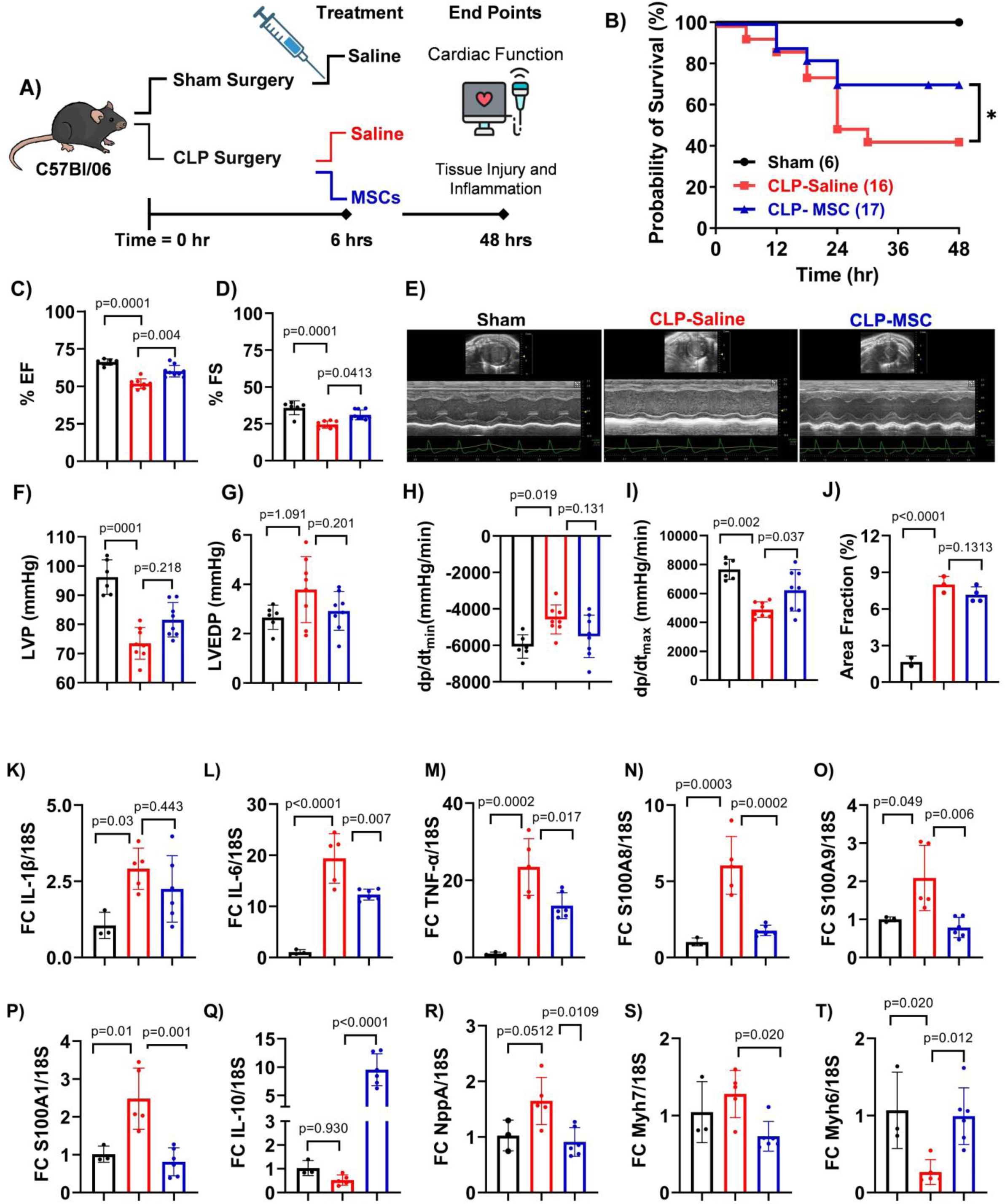
**Human bone marrow-derived MSCs (MSCs) improve survival, cardiac function, and reduce inflammation in a murine model of sepsis**. Schematic of experimental design (A). Male C57BL/6 mice randomized to either sham or cecal ligation and puncture (CLP) were further randomized at 6 hours post-surgery to receive intravenous MSCs (2.5×10⁵ cells/mouse) or equal volume saline. All mice were treated with subcutaneous fluid resuscitation (30 mL/kg), buprenorphine (0.1 mg/kg), and imipenem (25 mg/kg) every 8 hrs. Mice were euthanized 48 hours post-surgery for functional and molecular analyses. Kaplan–Meier survival analysis over 48 hours (CLP-MSC, n=17; CLP-saline, n=16; Sham, n=6, log-rank Mantel–Cox test) (B). Echocardiographic quantification of left ventricular ejection fraction (LVEF) (C) and fractional shortening (FS) (D). Representative M-mode echocardiograms from Sham, CLP-saline, and CLP-MSC groups (E). Invasive hemodynamic measurements; left ventricular (LV) pressure (F), LV end-diastolic pressure (G), dp/dt_min_ (H), and dp/dt_max_ (I). Histological assessment of myocardial polymorphonuclear (PMN) infiltration quantified as number of PMNs in a random area and shown as % area fraction (J). Fold change in LV gene expression (normalized to 18S) for pro-inflammatory markers IL-1β (K), IL-6 (L), TNF-α (M); damage associated molecular pattern receptor ligands S100A8 (N), S100A9 (O), S100A1 (P); anti-inflammatory cytokine IL-10 (Q); heart failure markers natriuretic peptide A (NppA) (R), β-myosin heavy chain (Myh7) (S), α-myosin heavy chain (Myh6) (T). Data are presented as mean ± SEM. ANOVA followed by Bonferroni-Dunn post hoc test were used, n=3-8/group.

MSC administration resulted in a trend towards decreased LV interleukin 1 beta (IL-1β, Figure 1K) expression, that did not reach significance, but a significant decrease in IL-6 (Fig 1L), tumor necrosis factor alpha (TNFα, Figure 1M), and cardiac alarmins ligands S100A8 (Figure 1N), S100A9 (Figure 1O), and S100A1 (Figure 1P). In keeping with previous data,[13] MSCs administration resulted in a ∼10-fold increase in the expression of anti-inflammatory cytokine IL-10 (Figure 1Q). MSCs also resulted in a decrease in the expression of hypertrophic phenotype markers: natriuretic peptide precursor A (NppA, Figure 1R) and fetal gene beta myosin heavy chain (βMhc, Myh7, Figure 1S) with a reconstitution of alpha Mhc (αMhc, Myh6, Fig 1T) levels. Taken together, this data supported the central thesis that IV MSC administration prevented the synthesis of myocardial DAMPs, sepsis-induced myocardial organ dysfunction, mortality, and reactivation of the fetal gene transcriptional program in septic hearts. The observed dissociation between IL-1β and IL-6 responses may reflect the timing of measurement or suggest MSC-mediated regulation of alternate signaling pathways upstream of IL-6 production.

### MSC-EV-mediated inhibition of inflammatory gene expression in cardiomyocytes

MSC-derived EVs are proposed to mediate paracrine anti-inflammatory activity. To determine whether EVs derived from MSCs reduced the expression of pro-inflammatory genes in cardiomyocytes, we isolated EVs from human bone marrow-derived colony-expanded and immortalized MSCs (ciMSCs), and compared them to EVs derived from platelets (Plt-EVs). Both ciMSC-EVs and Plt-EVs were prepared and characterized according to published criteria (MISEV2023, supplemental Figure 1A-C).[14] Primary murine neonatal cardiomyocytes were isolated and exposed to lipopolysaccharide (LPS), a key Gram-negative endotoxin that drives pro-inflammatory gene expression (supplemental Figure 1D). ciMSC-EVs, but not Plt-EVs, significantly reduced LPS-induced upregulation of IL-6 and IL-1β mRNA in cardiomyocytes. The protective effect of ciMSC-EVs was blocked by Dynasore, a small-molecule inhibitor of dynamin that prevents the endocytosis of EVs, suggesting that EV uptake is required for their anti-inflammatory effects (supplemental Figure 1E and F).

### IV MSC-EVs improved myocardial function and conferred survival benefits to septic mice

We next administered ciMSC-derived EVs to septic mice. In this experiment, both female and male C57Bl/6J mice, were randomized to CLP or sham surgery, and further randomized to receive IV equal volume saline, EVs-Plts, or ciMSC-EVs corresponding to cell equivalent of 6.45 × 10^8^ cells per mouse. Again, all mice received anti-biotics, fluids and pain management. EVs were administered 6 hours post-surgery, consistent with the MSC treatment protocol (schematic is presented in Figure 2A and protocol in detailed material and methods).

**Figure 2.**
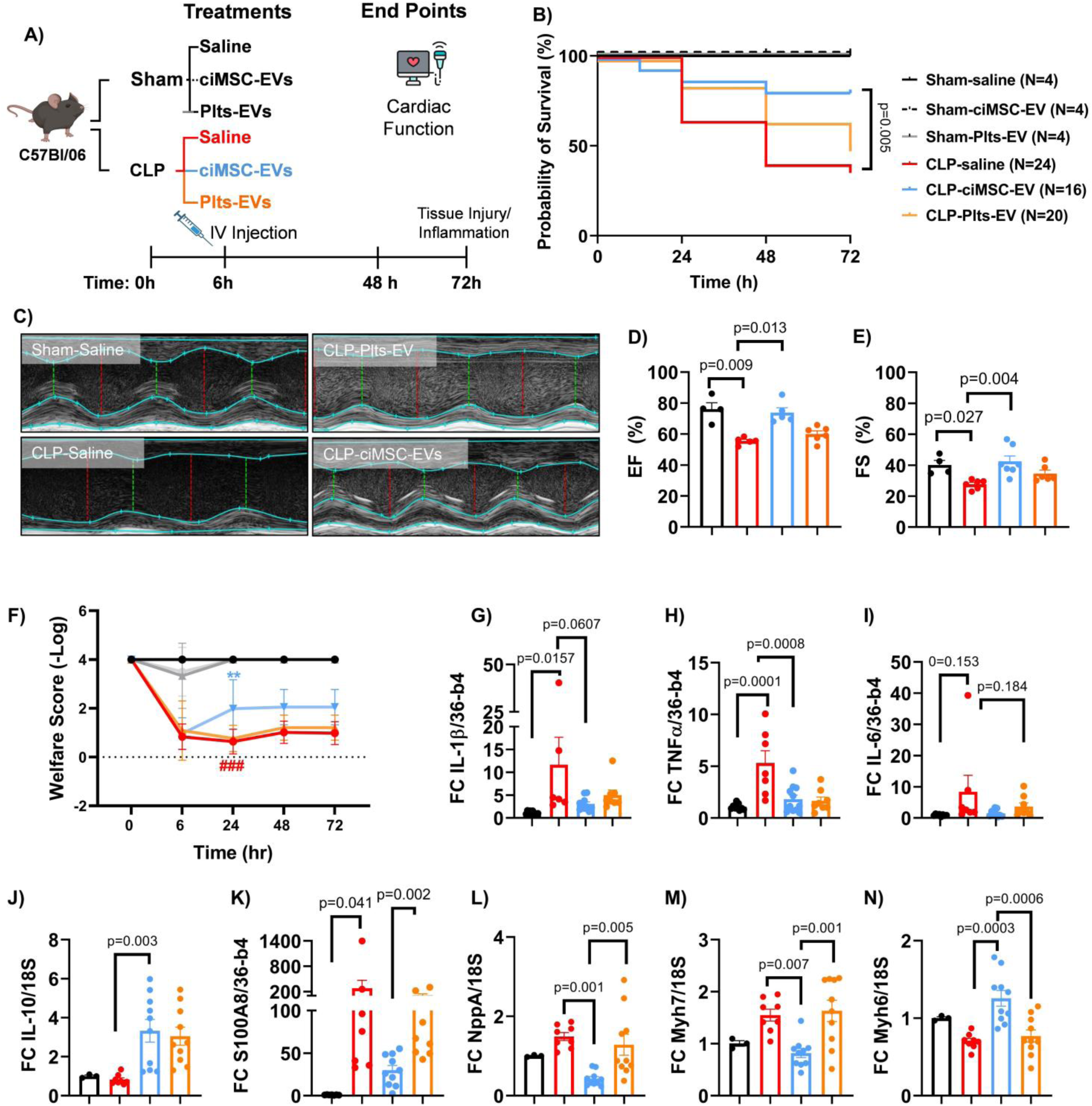
Extracellular vesicles derived from human ciMSCs, but not platelet-rich plasma, improve cardiac function and decreased mortality in septic mice. Experimental design and treatment scheme (A). Male C57BL/6 mice were randomized to sham or cecal ligation and puncture (CLP) surgery. At 6 hours post-surgery, mice received intravenous saline, clonally expanded and immortalized mesenchymal stem cell (ciMSCs) extracellular vesicles (EVs), or platelets (Plt)-EVs (2.5×10⁸ particles/mouse). All mice received standard supportive care (fluids, analgesia, antibiotics). Mice were monitored for 72 hours. Kaplan–Meier survival analysis across 72 hours (Sham-saline, n=4; Sham-ciMSC-EV, n=4; Sham-Plts-EV, n=4; CLP-saline, n=24; CLP-ciMSC-EV, n=16; CLP-Plts-EV, n=20) (B). Representative M-mode echocardiograms at 72 hours post-CLP (C), with quantification of % left ventricular ejection fraction (EF) (D) and fractional shortening (FS) (E) shown as mean±SEM. Welfare scores (a measure of pain severity and mobility where 1 = severe pain and low mobility to 4 = no pain and normal mobility) in groups as above across timepoints post-CLP (F). Fold change in cardiac mRNA levels (normalized to 18S) for pro-inflammatory markers IL-1β (G), TNF-α (H), IL-6 (I); anti-inflammatory cytokine IL-10 (J); damage associated molecular pattern receptor ligand S100A8 (K). Fold natriuretic peptide A (NppA) (L), β-myosin heavy chain (Myh7) (M), α-myosin heavy chain (Myh6) (N). Data are presented as mean±SEM. ANOVA followed by Bonferroni-Dunn post hoc test was used. n=3–10/group.

Seventy-two hours post-EV IV administration, CLP-mice that received ciMSC-EVs had an 81% survival rate compared to 36% in saline (antibiotics alone)-and 33% in Plt-EV–treated mice (Figure 2B). ciMSC-EVs enhanced LVEF by 32% and FS by 56% compared to saline placebo (Figure 2C-E). ciMSC-EVs resulted in a significant improvement in welfare scores (Figure 2F). In keeping with an anti-inflammatory effect, both ciMSC-and Plt-EVs resulted in decreased LV expression levels of genes IL-1β (Figure 2G), TNFα (Figure 2H), IL-6 (Figure 2I), and increased IL-10 (Figure 2J). Only mice that received EVs derived from ciMSCs demonstrated a decrease in the expression of S100A8 (Figure 2K), NppA (Figure 2L), Myh7 (βMhc, Figure 2M) and increase in Myh6 (αMhc) mRNA levels (Figure 2N).

In an independent set of experiments, blood chemistry analysis of end-organ damage at 72 hours post-CLP showed that mice treated with ciMSC-EVs had a consistent trend of improved absolute values of organ dysfunction, but these did not reach statistical significance (Supplemental Table 1). Of clinical relevance, while only ciMSC-EVs improved overall mouse welfare and mortality, PMN infiltration was significantly decreased in both groups (Supplemental Figure 2A and B). Histopathological analyses of non-cardiac organs showed that both EV types attenuated sepsis-induced liver and lung injury. In the liver, ciMSC-EV reduced both hepatic necrosis and steatosis relative to CLP-saline mice (Supplemental Figure 2C-E). Similar findings were found in lungs, where ciMSC-and Plt EVs treated mice exhibited reduced neutrophil infiltration compared to CLP-saline controls, with preservation of standard alveolar architecture (Supplemental Figure 2F-G), suggesting both types of EVs can dampen chemokine gradient and reduce neutrophil traffic.

### EV-miR sequencing and identification of miR-187

We hypothesized that post-transcriptional regulatory miRs, contained in MSC-EVs, may contribute to the survival benefit imparted by MSC-EV-administration in sepsis. To investigate this and to determine differences in miRs composition in MSC-EVs compared to control Plt-EVs, we sequenced miRs contained in both EV types. A total of 82 miRNAs were significantly enriched (fold change ≥2, FDR ≤ 0.05) in ciMSC-EVs compared to Plt-EVs (Figure 3A-C, supplemental Figure 3A), including miR-187a-3p (hereafter referred to as miR-187). This finding was in keeping with previous research from our group showing MSC administration enhanced the expression levels of endogenous miR-187 and its putative targets in septic hearts. In that study, increased endogenous miR-187 levels were associated with improved survival.[7] We next determined that cardiac miR-187 levels were reduced in murine hearts from CLP saline-treated compared to ciMSC-treated mice (Figure 3D). Similarly, miR-187 levels were increased after treatment with ci-MSC-EVs compared to saline or control Plt-EVs (Figure 3E). DIANA-TarBase v9,[15–17] identified 23 experimentally supported miR-187-targets, including TNFα and S100A4 (supplemental Figure 3B). Unexpectedly, IntaRNA[18–20], predicted the binding of miR-187 (supplemental Figure 3C), but not a scrambled control (supplemental Figure 3D) to the 3’ untranslated region (UTR) of the mRNA for IL-6 (hybridization energy-18.9 kcal/mol). Transfection of the miR-187 scrambled sequence (Figure 3F) failed to inhibit the luciferase activity from the IL-6 3’UTR reporter construct following treatment with LPS, whereas binding of miR-187 to the wild type IL-6–3’UTR inhibited luciferase activity, confirming binding of miR-187 to the 3’UTR of 1L-6 (Figure 3F), indicating IL-6 is a newly identified target of miR-187.

**Figure 3.**
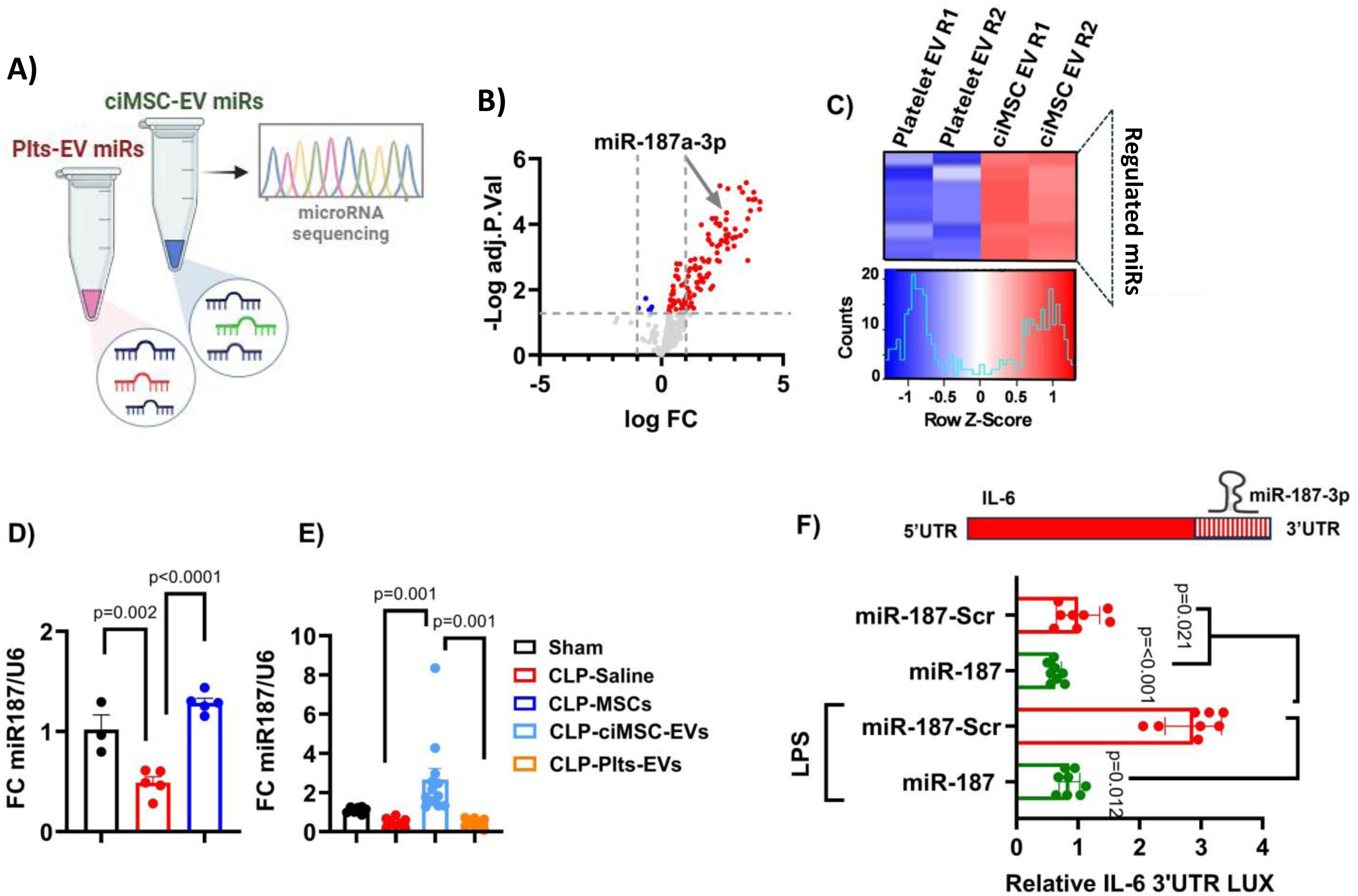
miR-187 is selectively enriched in ciMSC-EVs and regulates IL-6. Schematic highlighting sequencing of microRNAs isolated from clonally expanded and immortalized mesenchymal stem cell (ciMSCs) extracellular vesicles (EVs), vs. platelets (Plts) EVs (A). Volcano plot of differentially expressed miRs in ciMSCs-EVs vs. Plts-EVs with red dots indicating miRs overrepresented with p<0.05 highlighting miR-187a-3p (miR-187), blue dots log-fold enrichment greater than Log_2_=1 and grey p>0.05 (B). Heatmap of top enriched miRNAs in ciMSC-EVs vs. Plt-EVs including batch replicates (C). Fold-change in cardiac miR-187 normalized to U6 in sham, CLP-saline, and CLP–MSC groups (D), and in sham, CLP-saline, CLP-MSC, CLP-ciMSCs, and CLP–Plt-EV groups (E), presented as mean±SEM. Validation of miR-187 binding to the 3’ untranslated region (UTR) of the mRNA for IL-6 (F). Neonatal cardiac myocytes were co-transfected with the IL-6–3’UTR luciferase reporter and either miR-187 mimic or scrambled (SCR) miR and treated with lipopolysaccharide (LPS) (10 μM) or vehicle (PBS) for 24 h. Bars represent mean±SEM of luciferase expression presented as relative light units normalized to vehicle-treated cells. *p<0.05 vs vehicle control. n=3 (each in triplicate).

### miR-187 reduced inflammation and enhanced cardiac genes expression in cardiomyocytes

To determine if miR-187 regulates cardiomyocyte inflammation during sepsis, primary neonatal cardiomyocytes were isolated from 1–3-day-old CD1 mice and transfected with either a miR-187-3p mimic (miR-187) or a scrambled miRNA (negative control, NC). A schematic of gain-of-function experiments is shown in Figure 4. miR-187 delivery was confirmed by qRT-PCR before RNA sequencing (supplemental Figure 3E). NC and miR-187 transfected cells were then exposed to LPS or saline (24 hours). Total RNA was isolated and sequenced (see detailed methods).[7] Differential gene expression analysis identified 266 genes that were increased in response to LPS (LPS/NC) and decreased after miR-187 transfection (LPS+MIM/LPS), compared to NC (FDR≤0.05, FC≥Log2, Figure 4B). LPS-regulated genes whose expression was decreased by miR-187 transfection were primarily involved in TNFα, NF-KB and IL-6/JAK/STAT3 signalling (gene set enrichment analysis, GSEA) (Figure 4C). Pathview was used to map miR-187-treatment-responsive genes (Figure 4D) to NF-KB components, adhesion molecules (Vcam, Icam, and Selp),[21] chemokines (Ccl2, Ccl5, Cxcl2 and Cxcl3), cytokines (IL-18, TNFα, and IL-27) and coagulation/thrombosis factors (Plau, F10, F2R and FIP2) involved in cardiomyocyte pro-inflammatory pathway.[21, 22] Changes in the expression levels of key genes, including miR-187a-3p (Figure 4H), were validated in cardiomyocytes in separate experiments (Figure 4). Compared to LPS alone, miR-187 delivery reduced expression levels of TNFα (70%, Figure 4I), IL-6 (69%, Figure 4J), IL-1β (95%, Figure 4K).

**Figure 4.**
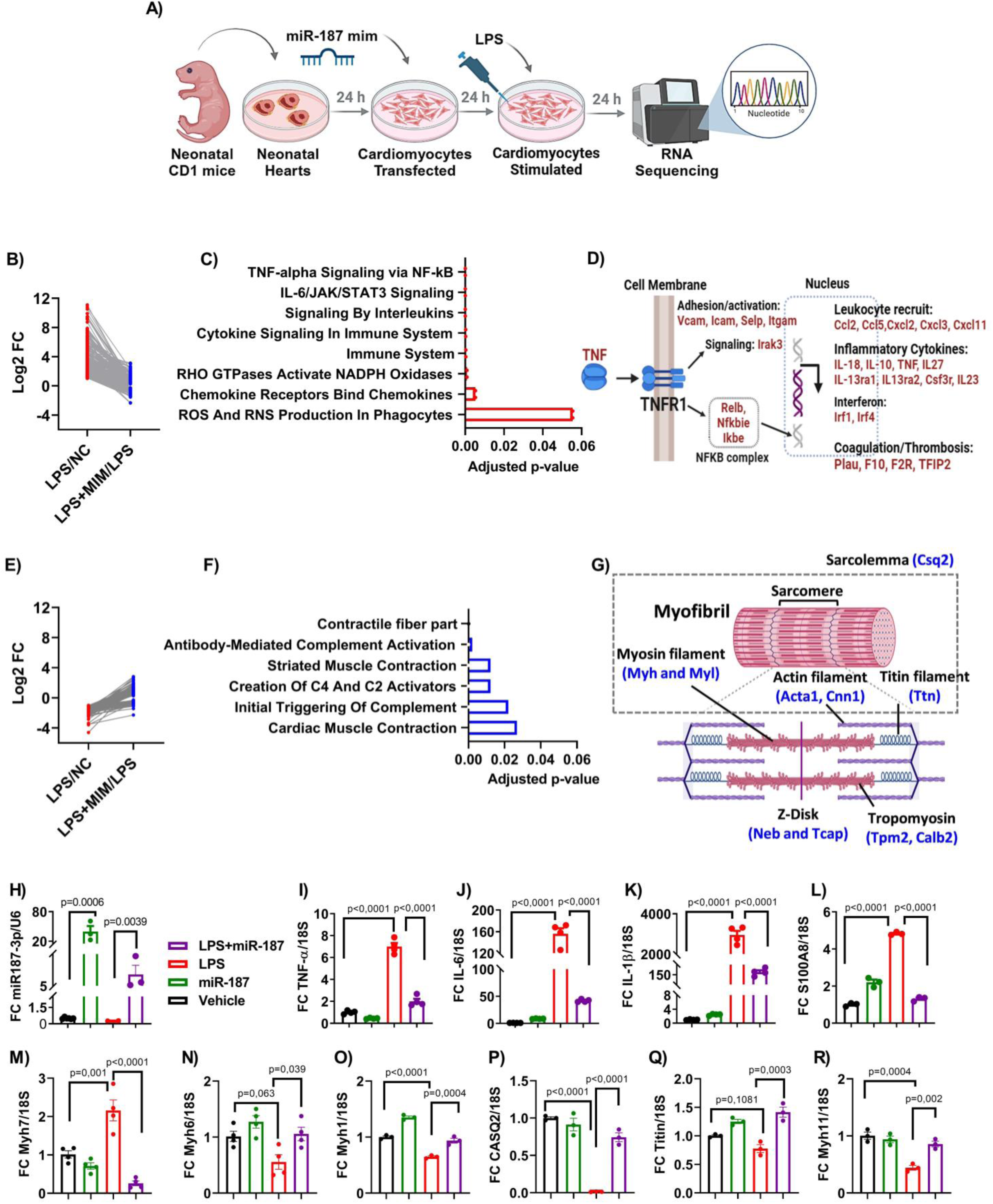
miR-187-3p mimic attenuates inflammatory signaling and restores cardiac gene programs in LPS-stimulated neonatal cardiomyocytes. Schematic of experimental workflow: neonatal mouse hearts were harvested, dissociated cardiomyocytes were transfected with miR-187-3p mimic or scrambled control, stimulated with LPS (1 µg/mL) for 24 hrs, and subjected to bulk RNA sequencing (A). Differential gene expression comparing LPS vs. LPS+miR-187 mimic treatment, showing genes downregulated in the mimic group (B), with pathway enrichment analysis of downregulated transcripts highlighting suppression of TNF-α/NF-κB and IL-6/STAT3 signaling (C). Schematic overview of TNFR1-mediated inflammatory signaling suppressed in mimic-treated cells (D). Genes upregulated in the miR-187-3p mimic group (E), and pathway enrichment indicating restoration of cardiac contraction, sarcomere structure, and complement regulation pathways (F). Schematic showing sarcomere organization and genes enriched in mimic-treated cells, including actomyosin, titin, troponin, and Z-disk components (G). Data reflect log2 fold change and Benjamini-Hochberg–adjusted p-values for gene set enrichment. In separate experiments, neonatal cardiomyocytes were transfected with miR-187-3p mimic (10 nM) or negative control (vehicle, transfection reagent only) for 24 hours, then stimulated with LPS (1 µg/mL) or vehicle (PBS) for an additional 24 hours. Fold change(s) of miR-187 normalized to U6 (H); TNF-α (I), IL-6 (J), IL-1β (K), S100A8 (L), β-myosin heavy chain (Myh7) (M), a-myosin heavy chain (Myh6) (N), Myh1 (O), calsequestrin 1(Casq2) (P), Titin (Q), and Myh11 (R) all normalized to 18S and presented as mean±SEM. One-way ANOVA with Bonferroni-Dunn post hoc test was used for statistical comparisons. n=3-4/group.

In contrast, miR-187 transfection resulted in increased mRNA levels (LPS+MIM/LPS) of 174 genes that were otherwise decreased in response to LPS (LPS/NC, FDR≤0.05, FC≥Log2, Figure 4E-G). Genes responsive to miR-187 encoded for contractile fiber and filament protein components including myosin heavy (Myh) and light (Myl) chains, actin (Acta1, Cnn1), and Titin (Ttn). In addition, contraction-related proteins such as calsequestrin (CSq2/CASQ2), tropomyosin (Tpm2 and Calb2), and Z-disk-related proteins (Neb and Tcap). In independent experiments, transfection of miR-187 led to decrease in the expression levels of S100A8 (80%, Figure 4L), heart failure marker βMhc (myh7, Figure 4M) and reconstitution of mRNA levels for cardiac structural and contractile proteins αMhc/Myh6 (100%, Figure 4N), Myh1 (33%, Figure 4O), Casq2 (500%, Figure 4P), Titin (133%, Figure 4Q) and Myh11 (50%, Figure 4R). Analysis of miR-187 regulated genes revealed they did not all share miR-187 regulatory sequences.

An alternative explanation for coordinated transcription of cardiomyocyte-related genes may be co-regulation of cardiac-specific transcription factors that in turn coordinate the expression of these functionally related genes. Supplemental Table 4A shows co-regulation of top transcription factors MyoD (myogenic differentiation 1), Tbx15 (T-Box Transcription Factor TBX15) and Mef2C (Myocyte Enhancer Factor 2C) associated with increased expression of the 174 cardiac-related genes in septic cardiomyocytes transfected with miR-187.[23] IntaRNA predicts binding of miR-187 to 3’UTR of both human and murine MyoD, Tbx15 [24] and Mef2C genes (supplemental Table 4B source files). In vivo validation of miR-187-regulation of cardiac transcription factors was performed in subsequent experiments.

### Preparation and cellular effect of Lipid Nanoparticles for miR-187 delivery

To deliver miR-187 in vivo to septic mice, we generated lipid nanoparticles (LNPs) encapsulating a proprietarily modified miR-187 mimic (patent under revision ref: 579-022-P). Our LNPs were comprised of an ionizable lipid (DLin-MC3-DMA), cholesterol, a structural helper lipid distearoylphosphatidylcholine (DSPC), and PEGylated lipid (DMG-PEG2000, Figure 5A). Standardized microfluidics (NanoAssemblr/Precision Nanosystems) was used to mix the lipid-and miR-drug-containing solutions. Standardized characterization methods were used to confirm the quality of LNP-miR-187 formulation: diameter (∼113 nm) and polydispersity index (≤0.3, PDI) (supplemental Figure 4A), encapsulation efficiency (>95% supplemental Figure 4B) and Z-potential (ZP, ∼-2.4mV, supplemental Figure 4C). LNP structural integrity was also assessed by electron microscopy (Figure 5B).

**Figure 5.**
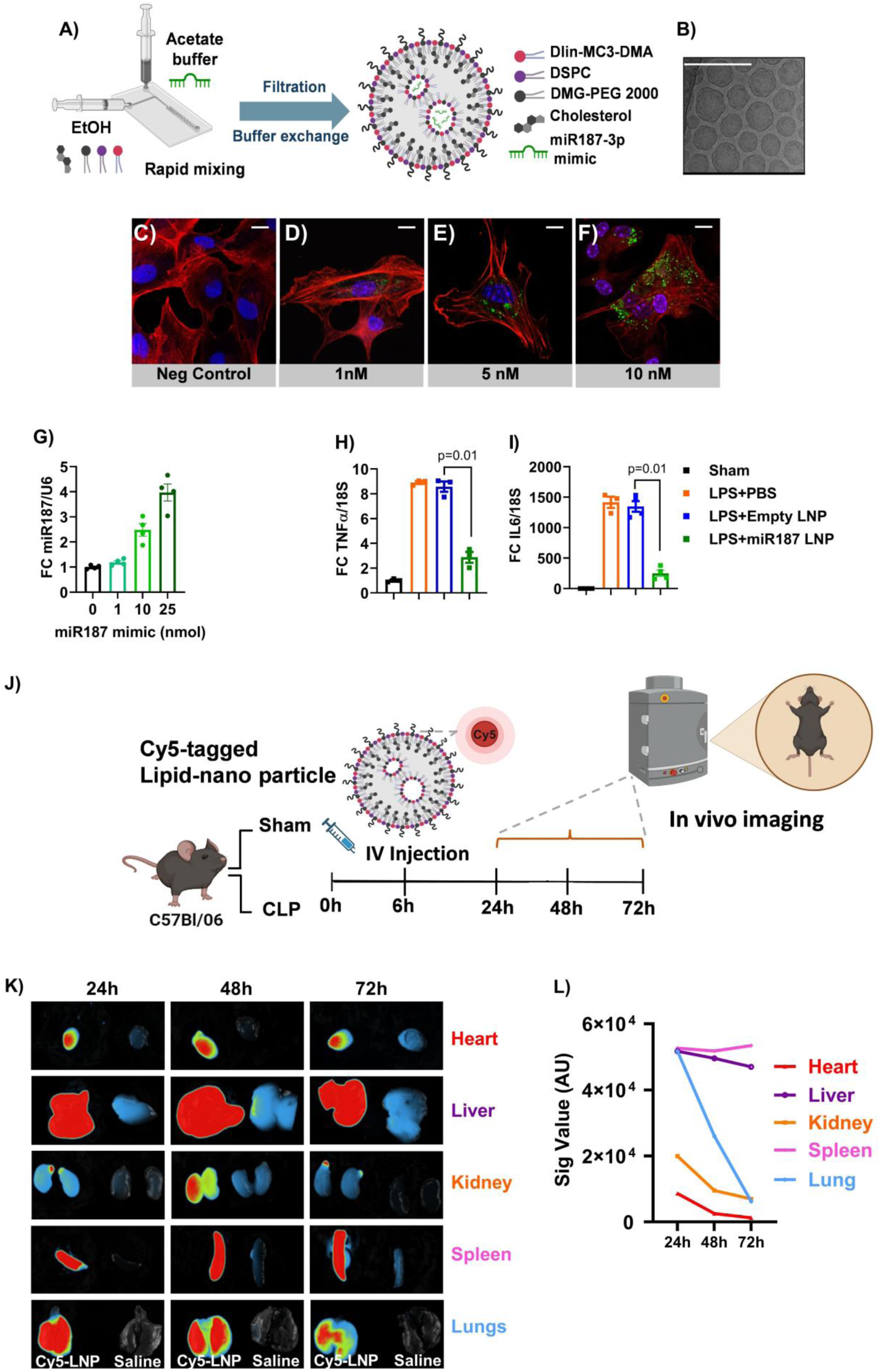
miR-187-3p mimic-loaded lipid nanoparticles (LNPs) are up taken by cardiomyocytes and septic murine hearts. Schematic of LNP formulation by rapid ethanol injection method, incorporating ionizable lipid (Dlin-MC3-DMA), DSPC, cholesterol, PEG-lipid, and either control or miR-187-3p mimic (A). Transmission electron microscopy (TEM) images of LNPs loaded with miR-187-3p mimic (B). Confocal images of neonatal cardiomyocytes uptake of unlabelled (negative control) (C) or fluorescently labelled miR-187-3p mimic LNPs at increasing doses (1, 5, and 10 nM; panels C-F), stained for actin (red), nuclei (blue), and LNP uptake (green). Fold-change in cellular miR-187 normalized to U6 (G) and impact on expression levels of TNFα and IL-6 normalized to 18S after treatment with LPS (1 µg/mL) or vehicle (PBS) for 24 hours. One-way ANOVA with Bonferroni-Dunn post hoc test was used for statistical comparisons. n=3-4/group. Schematic of in vivo imaging workflow: C57BL/6 mice underwent sham or cecal ligation and perforation (CLP) surgery, followed by intravenous injection of Cy5-labeled miR-187 LNP or saline at 6 hours post-surgery and serial imaging up to 72 hours (J). Ex vivo imaging of organs (heart, liver, kidney, spleen, lungs) at Day(s) 1, 2, and 3 post-injection (K), with quantification of signal as above (L). n=1-2/group.

Primary murine neonatal cardiomyocytes were exposed to increasing amounts (0, 1, 5, and 10 nM) of fluorescently tagged LNP-miR-187 (Figures 5C–F). Fluorescence microscopy confirmed a dose-dependent increase in LNP accumulation, with the highest uptake at 10 nM (Figure 5F). We confirmed delivery of miR-187 (payload) at this concentration by qRT-PCR (Figure 5G); this concentration of 10nM was used for follow-up in-vitro studies to show target effect, determined as a decrease in mRNA levels of target genes TNFα (Figure 5H) and IL-6 (Figure 5I) and cytotoxicity studies. Neonatal cardiomyocytes or cells (AC10) derived from primary human adult ventricular tissue were exposed to increasing concentrations of the therapeutic LNP-miR-187 formulation. Absence of cytotoxicity was determined by measuring cell viability (Supplemental Figure 4D), lactate dehydrogenase release (LDH, Supplemental Figure 4E), and percent early and late apoptosis (Supplemental Figure 4F) after 24 hrs exposure. Proliferation assay (BrdU, or 5-bromo-2’-deoxyuridine) demonstrated LNP-miR-187 does not lead to increased cellular proliferation (Supplemental Figure 4G). Cells were also transfected with a Nuclear Factor Kappa Beta (NFκβ) reporter construct and exposed to LNP-miR-187 to demonstrate the therapeutic formulation does not lead to NFκβ activation (Supplemental Figure 4H).

### In vivo biodistribution of LNP-miR-187 formulation after intravenous delivery

Before evaluating the therapeutic effects of LNP–miR-187 in vivo, we determined its biodistribution following IV injection. Fluorescently tagged LNPs (1 nmol miRNA/mouse) were injected in sham or CLP mice and tracked over time (Supplemental Figure 4G). Daily imaging over 14 days showed progressive decrease in fluorescent signal, with near-complete clearance by day12. In a separate and independent cohort, LNP-miR-187 nanoparticles were injected 6 hr post CLP and tracked over the first 3 days. Tagged LNP-miR187 could be detected in various sepsis relevant organs including the liver, spleen, lungs, and with lower but detectable and quantifiable levels in the heart and kidneys at 24-, 48-, and 72-hours post-injection (Figures 5K-L).

### LNP-miR-187 mitigated cardiac dysfunction and enhanced probability of survival in septic mice

Equal numbers of female and male C57BL/6 mice (12–14 weeks) were randomized to CLP or sham surgery and, 6 hours post-surgery, were further re-randomized to receive IV LNPs (6.3 × 10¹¹ particles/mouse) containing either scrambled miR (SCR negative control, 1 nmol/mouse), miR-187 mimic (LNP-miR-187, 1 nmol/mouse), or saline (Figure 6A). In antibiotic-treated and fluid-resuscitated mice, the early probability of survival at 48 hours markedly improved in LNP-miR-187–treated mice compared to controls from 48 to 100% (Figure 6B). Animal welfare scores also improved 6 hours post-CLP (6 hours after IV delivery of LNP–miR-187, Figure 6C).

**Figure 6.**
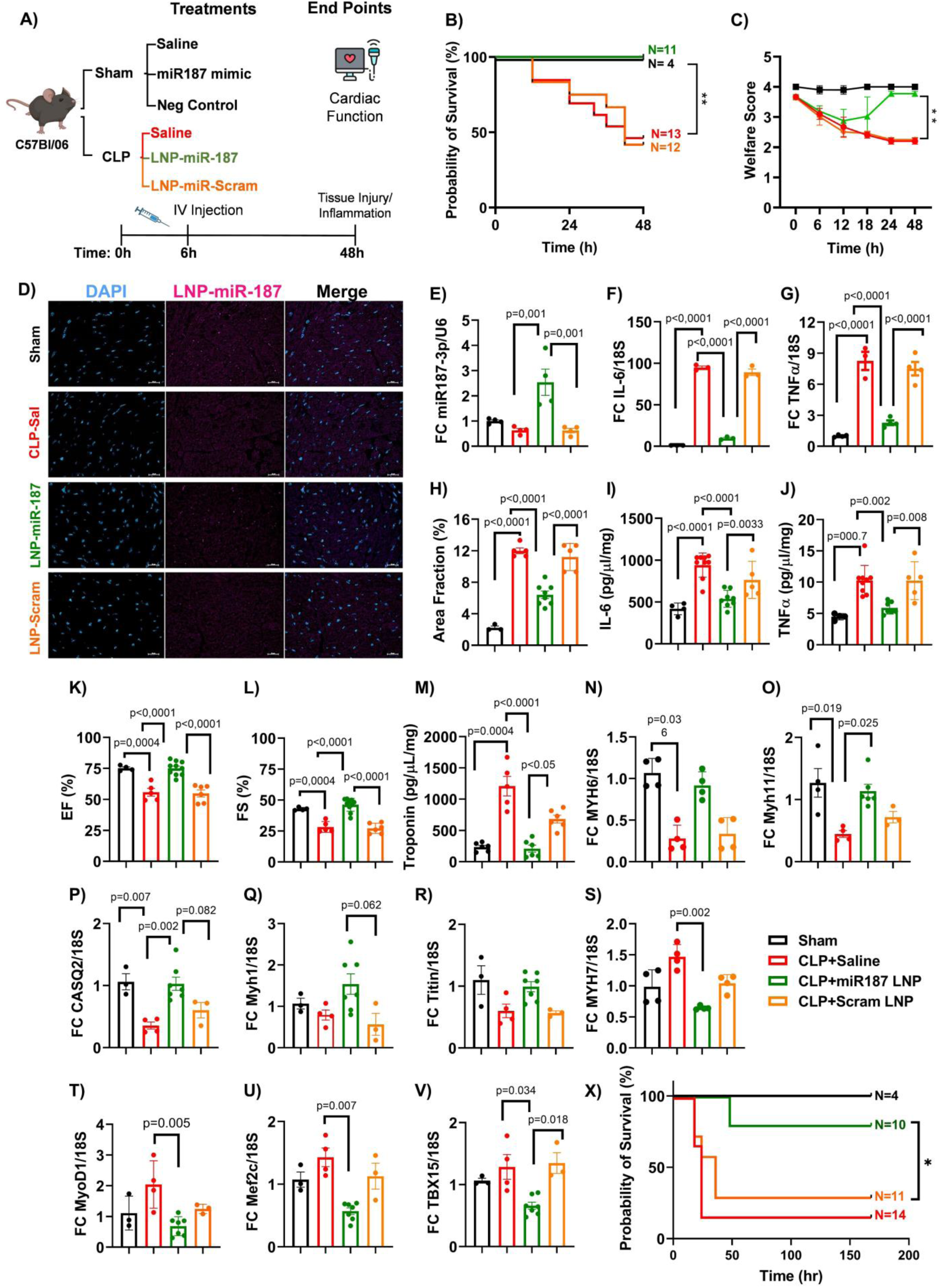
miR-187-3p mimic-loaded LNPs improve survival, cardiac function, and reduce inflammation in septic mice. Schematic of experimental design: C57BL/6 mice underwent cecal ligation and perforation (CLP) or sham surgery and were randomized at 6 hours post-surgery to receive saline, miR-187 mimic LNPs (10 µM), or scrambled (SCR) LNPs via intravenous injection and followed for 48 hours (A). Kaplan–Meier survival analysis over 48 hours showing % probability of survival (B), and serial welfare scores (a measure of pain severity and mobility where 1 = severe pain and low mobility to 4 = no pain and normal mobility) (C) (**p<0.01, n=6-15/group). Representative miRscope in situ hybridization images showing cardiac expression of miR-187 (magenta) and DAPI (blue) and merged image in sham, CLP-saline, CLP+miR-187 LNP, and CLP+negative control LNP (D) and relative cardiac miR-187 normalized to U6 (E). Relative IL-6 (F), TNF-α (G), mRNA normalized to 18S. Histological quantification of myocardial lymphocyte infiltration reported as number of lymphocytes in a random area and shown as % area fraction (H). LV tissue levels of IL-6 (I), TNF-α (J) protein levels (ng/mL/mg tissue). Left ventricular ejection fraction (EF) (K), fractional shortening (FS) (L) shown as percentage and cardiac troponin (M) (pg/mL tissue lysate). Myh6 (M), Myh11 (N), Casq2 (O), Myh1 (P), Titin (Q), Myh7 (R), Mef2c (S), and Tbx15 (T), all normalized to 18S. Data are presented as mean±SEM, n=5-15/group. Kaplan– Meier survival analysis showing % probability of survival over 7 days (U). Statistical comparisons were performed using one-way ANOVA with Bonferroni-Dunn post hoc test or log-rank test for survival.

Blood chemistry showed improved organ function, including reduced circulating troponin, lactate, and liver enzyme levels following miR-187 delivery (Supplemental Table 2). miRNAscope showed treatment with miR-187-LNP enhanced miR-187 punctae in cardiac tissue 2.5-fold compared to scrambled control (Figure 6D-E) and decreased LV expression of IL-6 (Figure 6F) and TNFα (Figure 6G). Decrease in inflammatory gene expression was associated with a decrease in LV leukocyte infiltration (Figure 6H), IL-6 and TNFα protein levels (Figure 6I and 6J) in septic LVs. Decreased inflammation was also associated with an improvement in LVEF (Figure 6K), LVFS (Figure 6L) and decreased in plasma troponin levels (Figure 6M).

In parallel, intravenous LNP-miR-187 treatment resulted in increased Myh6 (Figure 6N), Myh11 (Figure 6O), Casq2 (Figure 6P), and Myh1 (Figure 6Q), with no effect on titin (Figure R), and decreased myh7 (Figure 6S). Myocardial gene reprograming (switch from alpha to beta Mhc) was associated with a decrease in the expression of known regulatory transcription factors Mef2c (Figure 6T), Txb15 (Figure 6U) and MyoD1 (Figure 6V). In an independent experiment, we extended the probability of survival study to 7 days and showed that treatment of CLP female and male mice with LNP-miR-187 improved the probability of survival by 70% (Figure 6W). We did not identify meaningful differences in survival probability between female and male mice.

Electron microscopy, demonstrated that LNP-miR-187 administration prevented sepsis-induced structural disruption of the LV contractile apparatus, as evidenced by preservation of Z-line distance (Supplemental Figure 5A and B). While having no effect of the number of mitochondria, treatment with LNP–miR-187 also prevented the development of mitochondrial shrinkage, a reduction in the size of mitochondria, a characteristic feature of mitochondrial dysfunction during sepsis (Supplemental Figure 5C). In human adult LV derived cardiomyocytes (A10) exposed to LPS, treatment with LNP-miR-187 markedly reduced expression of alarmins S100A8 and S100A9 genes compared to transfection of the miR-187 inhibitor (Supplemental Figure 6A and B).

### Safety and Systemic Tolerability of intravenous LNP-miR-187 in Healthy and Septic Mice

We administered escalating doses of LNP–miR-187 to healthy male mice (0.1, 1, and 3 nmol) alongside saline and LNP-scrambled controls and monitored mice for two weeks (Supplemental Table 2). LNP administration was well-tolerated at all tested doses. Blood chemistry profiles revealed no signs of organ toxicity, with liver enzymes, renal function, electrolytes, and serum proteins remaining within physiological ranges. A mild elevation in white blood cell counts was noted in all LNP-treated groups. Lymphocyte counts remained stable in the 1 nmol (therapeutic dose) group. Histological examination of key organs (lung, kidney, heart, brain, spleen, thymus, and intestine) revealed no pathological abnormalities across all treatment arms. A notable exception was vacuolization in liver sections of all LNP-treated mice (scrambled and miR-187 loaded), consistent with previously reported PEG-associated effects. These vacuoles were not associated with inflammation or tissue damage and are considered transient and non-pathogenic, as seen with other PEG-containing therapeutics like Patisiran and mRNA vaccines.[25]

We next evaluated the safety of LNP–miR-187 in the setting of sepsis. Serum biochemistry was assessed 48 hours post-treatment in CLP mice receiving saline, scrambled control LNP, or LNP– miR-187 and compared to sham-operated animals (Supplemental Table 3). CLP-saline mice showed biochemical evidence of multi-organ dysfunction, including markedly elevated AST (219.45 IU/L), low albumin (26 g/L), and altered lipid and glucose metabolism. In contrast, mice treated with LNP–miR-187 showed partial normalization of AST (195 IU/L) and higher albumin levels (24.15 g/L), suggesting hepatoprotective effects. Renal markers (creatinine, BUN) remained within normal ranges in all groups, and no significant electrolyte disturbances were observed. These results support the conclusion that LNP–miR-187 is systemically well-tolerated in both healthy and septic mice and may confer modest organ-protective effects under inflammatory stress.

### Co-delivery of MSC-EVs with LNP-miR-187 inhibitor or mimic did not significantly improve or abrogate the effects of MSC-EVs on sepsis survival

We investigated whether co-delivery of MSC-derived EVs and LNP-miR-187 mimic or inhibitor affected the effects of ciMSC-EV-treatment on the probability of survival 72 hrs following CLP (Figure 7A). IV ciMSC-EVs or LNP-miR-187 mimic (MIM) or inhibitor (INH) had no ill effect in sham mice (Figure 7B). The probability of survival in mice randomized to LNP-miR-187 was better than mice that received ciMSC-EVs. Co-administration of LNP-miR-187 and ciMSC-EVs did not improve the probability of survival from sepsis much beyond that seen with ciMSC-EV alone. Co-administration of the LNP-miR-187INH did not abrogate the beneficial effects of receiving the ciMSC-EVs. While both ciMSC-EVs and LNP-miR-187 reduced expression levels of the miR-187 target gene IL-6 (Figure 7C), LNP-miR-187INH resulted in marked decrease in the LV expression levels of cardiac specific genes Myh1 (Figure 7D), Myh11 (Figure 7E), Titin (Figure 7F) and Casq2 (Figure 7G). Expression levels of these cardiac specific genes in septic ventricles and in response to ciMSC-EVs and LNP-miR-187 was more varied and likely reflect heterogeneity in sepsis severity. LNP-miR-187 and ciMSC-EV administration attenuated the expression of cardiac transcription factors involved in LV sepsis-induced phenotypic switch MyoD (Figure 7G), Mef2c (Figure 7H), and Tbx15 (Figure 7I), suggesting a biological effect of miR-187 delivery in preventing sepsis-induced cardiac remodeling. We further evaluated the relationship between echo measures of LV dysfunction and LV miR-187 expression levels to demonstrate higher miR-187 levels are associated with improved LV function in mice that received ciMSC-EVs (Figure 7J) or miR-187 LNPs (Figure 7L). We calculated the treatment efficacy correlation ratio suggesting improved LV function associated with the increased miR-187 LV expression levels (Figure 7M).

**Figure 7.**
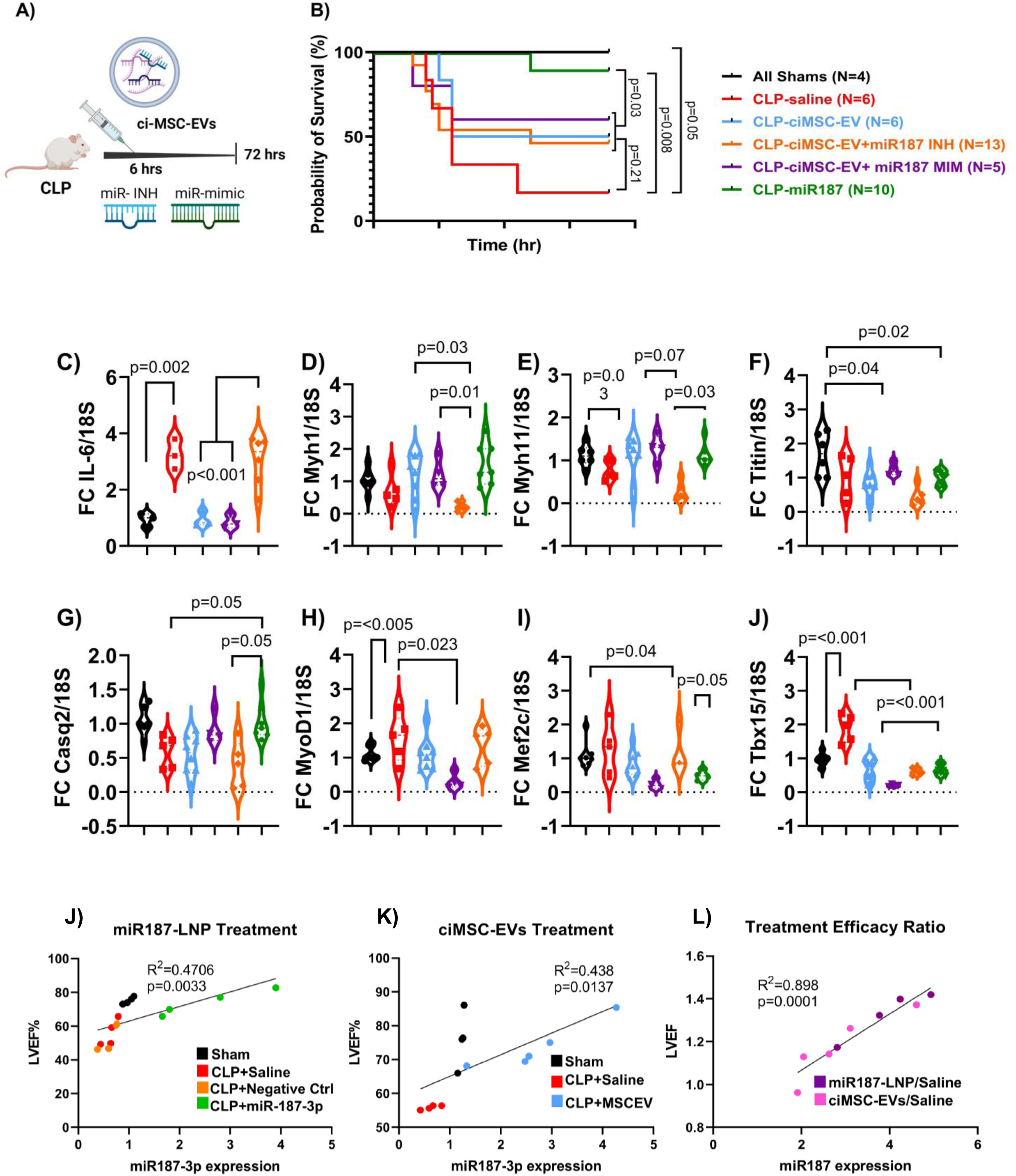
Co-delivery of ciMSC-EVs with LNP-miR-187 mimic or inhibitor does not significantly improve or abrogate the effects of ciMSC-EVs on sepsis survival. Experimental design and treatment scheme (A). Male and female C57BL/6 mice were randomized to sham (black) or cecal ligation and puncture (CLP) surgery. At 6 hours post-surgery, mice received intravenous saline (red), lipid nanoparticles (LNP) miR-187 formulation (green), or clonally expanded and immortalized mesenchymal stem cell (ciMSCs) extracellular vesicles (EVs, 2.5×10⁸ particles/mouse, blue) co-administered with (purple) or without (orange) lipid nanoparticles (LNPs) loaded with miR-187-3p mimic (miR-187 MIM) or miR-187-3p inhibitor (miR-187-INH). Four different sham animal groups were included (n=1/group) treated with either saline, ciMSC-EV, LNP-miR-187 MIM or LNP-miR-187-INH alone. None of the sham animals died and for simplicity are shown together (black line). All mice received standard supportive care (fluids, analgesia, antibiotics). Mice were monitored for 72 hours and Kaplan–Meier survival analysis performed (B). No difference in survival were noted between mice that received ciMSC-EVs with and without LNP-miR-187 MIM or LNP-miR-187-INH. To determine statistical significance in survival differences these were considered one group. Mice that received LNP-miR-187 MIM demonstrated a significant survival advantage over the first 72 hrs post CLP. While both ciMSC-EVs and LNP-miR-187 MIM nanoparticles were anti-inflammatory (decreased IL-6 C), administration of the LNP containing a miR-187-inhibitor consistently resulted in marked decrease in the LV expression levels of cardiac specific genes Myh1 (D), Myh11 (E), Tintin (F) and calsequestrin 1 (G). Expression levels of these cardiac specific genes in septic ventricles and in response to MSC-EVs and LNP-miR-187 was more varied and likely reflect heterogeneity in sepsis severity. Importantly however, LNP-miR-187 administration was the only treatment that resulted in decreased expression of cardiac transcription factors involved in left ventricular remodeling and reactivation of the foetal gene-program during sepsis MyoD (H), Mef2c (I) and TBX15 (J). Data represented as mean+SEM. n=4-8/group. Statistical comparisons were performed using one-way ANOVA with Bonferroni-Dunn post hoc test or log-rank test for survival.

### miR-187 levels in human sepsis and sepsis-induced myocardial dysfunction

We isolated microRNAs from formalin fixed paraffin embedded LVs from patients who died with sepsis (N=6) and compared to control patients who died with other non-septic and non-cardiac related problems (N=6); miR-187 copy number is significantly decreased in human septic hearts (Figure 8A). Using miRNAscope, we probed for miR-187 in situ in separate human cardiac autopsy samples and visually identified less miR-187 punctae in septic LVs (N=3) compared to non-septic (N=3, Figure 8B). Banked plasma samples from patients enrolled in our own observational clinical trial (INSIST), were used to show decreased miR-187 in circulating plasma-derived exosomes from septic (N=14) compared to critically ill non-septic patients (N=6, Figure 3C). In parallel, changes were accompanied by elevated plasma levels of IL-1β (Figure 3D) and TNFα (Figure 3E) in septic patients. To evaluate the clinical relevance of miR-187 levels in relationship to myocardial function in human sepsis, we quantified circulating miR-187 copy number in serum from patients with sepsis with or without echocardiographic evidence of presumed new myocardial dysfunction. Septic patients enrolled in the COLOBILI study with echocardiograph-proven preserved cardiac function (N=17, patient cohort is described in Supplemental Table 5) exhibited higher circulating miR-187 levels compared to patients with documented abnormal echocardiograms (N=13, Figure 8D, correlation r=0.45 p=0.009), suggesting that elevated miR-187 copy number is associated with a normal (>50%) EF as measured by echocardiogram. Pearson correlation matrix validated a correlation between miR-187 copy number with the sequential organ failure assessment score (p= 0.0015) and neutrophil to leukocyte ratio (p= 0.016) in patient with abnormal echocardiograms (supplemental Figure 7A), lactate levels (p=0.028) and mean arterial blood pressure (p= 0.037) in patient with no evidence of cardiac dysfunction (supplemental Figure 7B). Data is presented in supplemental Table 6.

**Figure 8.**
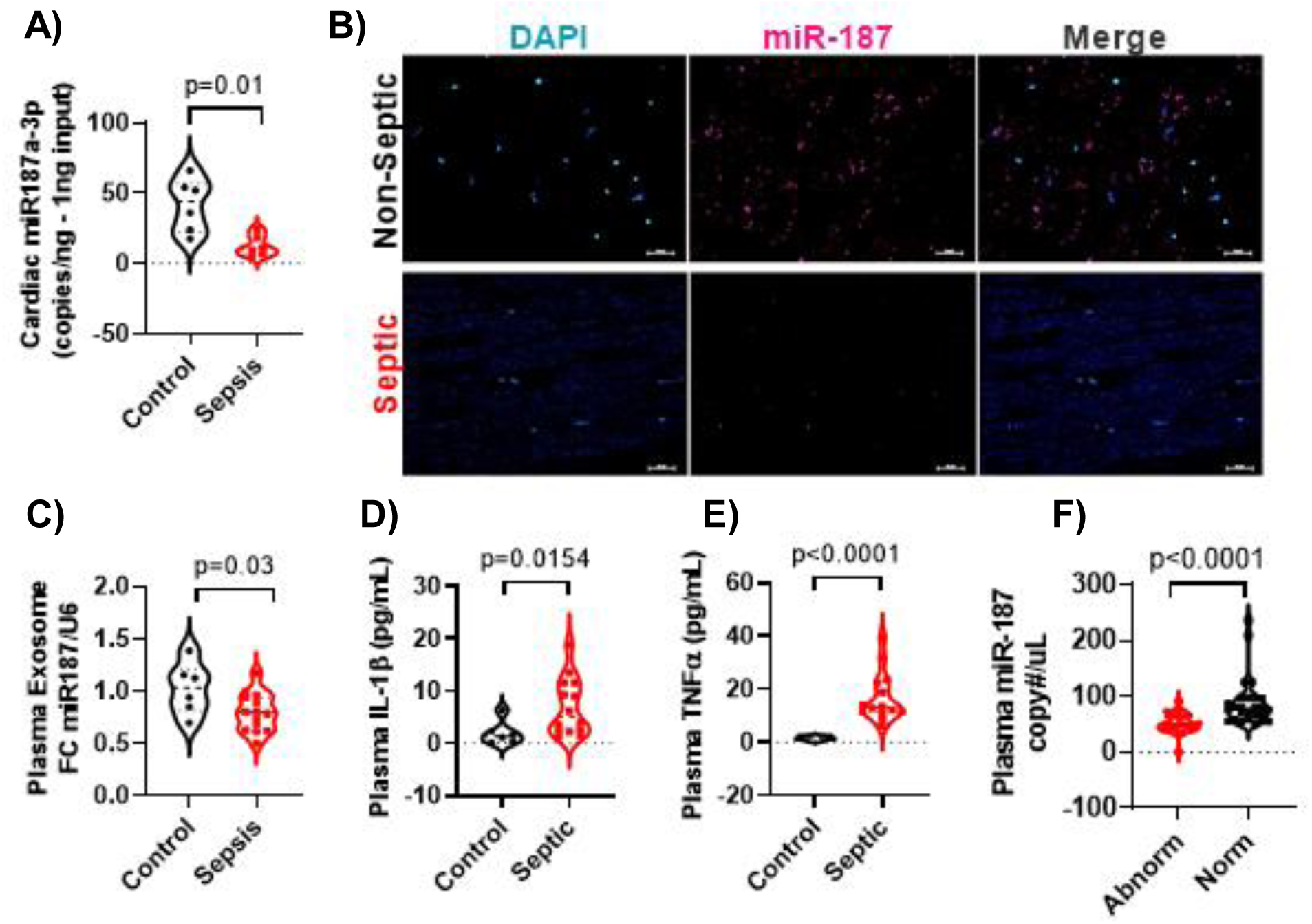
**Circulating miR-187 levels are altered in human sepsis and in patients with documented new septic-associated echocardiogram changes in cardiac function**. miR-187-3p copy number in post-mortem septic and non-septic cardiac tissue as analyzed by ddPCR (copies/ng input) (A). Representative miRscope assay images of miR-187-3p (magenta) in post-mortem human cardiac tissue from control (cancer) and sepsis patients (24–48 hrs post ICU admission); nuclei counterstained with DAPI (cyan) (B). Fold-change in miR-187 contained in exosomes isolated from plasma normalized to U6 (C), plasma IL-1β (D), TNFα (E), as analyzed by ELISA and shown as pg/mL. Plasma miR-187-3p copy number as determined by ddPCR in septic patients with normal vs. abnormal left ventricular ejection fraction (LVEF) (F). Data are shown as violin plots showing median and 25^th^ and 75^th^ percentile. n=7-46. The Wilcoxon Signed Rank Test was used for statistical comparisons.

To determine if in humans, circulating miR-187 levels are impacted by MSC treatment, we used microRNA isolated from PaxGene tubes collected during our Phase 1 clinical trial of Cellular Immunotherapy for Severe Sepsis (CISS) and measured circulating miR-187 copy-number. As anticipated, plasma miR-187 copy number was decreased in septic patients compared to healthy controls. In the 9 patients that received escalating doses of IV MSCs, we did not see a consistent dose-effect at baseline, 24 or 72 hrs after infusion; however, there was a small trend towards increased miR-187 copy number in samples collected 72 hrs post-MSC IV infusion (supplemental Figure 7C and D).

## Discussion

In this study, we demonstrate that systemic delivery of a novel miR-based therapeutic, LNP-miR-187, effectively mitigates cardiomyocyte and LV inflammation, prevents sepsis-induced heart-failure gene remodelling, improves LV ejection fraction and significantly reduces mortality in vitro murine, human cell models as well as in pre-clinical models of polymicrobial sepsis. Our findings offer compelling evidence in support for miR-mediated intervention in sepsis-induced myocardial dysfunction, for which currently there are limited therapeutic options and a substantial unmet clinical need. Unlike other approaches that may also have pleotropic effects, e.g. corticosteroids, vasopressors and/or inotropes, our approach leverages the role of miR-187 as a post-transcriptional regulator of translation to achieve systemic and tissue-specific, safe, gene modulation, paving the way for its potential translation into early-phase clinical trials. By integrating advances in microRNA chemistry with systemic delivery systems, such as lipid nanoparticle encapsulation, our study highlights the transformative potential of microRNA therapeutics to limit the contribution of LV dysfunction to sepsis-associated mortality.

Myocardial depression is a treatable trait of sepsis that is characterized by a cardiac output that fails to meet metabolic demands. Reduced LV ejection fraction occurs in 60% of patients during the first 3 days of treatment of septic shock.[26] Forty percent of these patients present with LV hypokinesia at admission, suggesting depression in LVEF can occur in the earliest stage of sepsis.

Approximately, 60% of patients who present with a clinical picture of cardiovascular impairment have increased mortality (from 70-90%), in contrast with 20% mortality in patients without cardiovascular involvement.[27] Approximately 15% of deaths related to septic shock are attributable to myocardial depression.[28, 29] In those patients who die, myocarditis is seen in 27% of autopsy samples as well as necrotizing vasculitis, bacterial colonization (11%), myocardial abscesses, necrosis of cardiac fibers (∼10%), and interstitial edema (28%) associated with degenerative myocardial changes suggesting the heart is a major target organ.[28, 29] LNP-miR-187 administration significantly attenuated LV dysfunction in our sepsis model and while a systemic effect cannot be excluded, given the experimental design, a specific effect of miR-187 on the regulation of cardiac-specific transcription factors that regulate reactivation of the foetal gene expression program strongly supports a cardiac-specific role of miR-187 in preventing the development of sepsis-associated heart failure.

We chose to deliver LNP-miR-187 systemically because sepsis is a systemic disease and while the liver is the primary location for the synthesis of TNFα during sepsis in vivo, LNP-miR-187 delivery reaches the heart, where it is detected for the first 72 hrs after administration and leads to decreased LV inflammation and cardiac gene remodelling. The lipid nanoparticles are Food and Drug Administration (FDA) approved and are currently in clinical use facilitating translation to future human studies. Our therapeutic nanoparticles are taken up by cardiomyocytes where they are able to attenuate inflammation and LPS-induced cardiac gene remodeling. Delivery of LNP-miR-187 did not affect cellular viability, LDH release, or cellular apoptosis. Proliferation studies suggested LNP-miR-187 reduced Phenyl and PMA induced cardiomyocyte proliferation. Importantly, LNP-miR-187 were not proinflammatory in either in vitro or in vivo. We also monitored the effects of LNP-miR-187 on histological, biochemical and haematological evidence of toxicity in healthy mice for 14 days and found no gross evidence of toxicity suggesting LNP-miR-187 delivery is safe. This is not surprising given that LNP-miR-187 delivery is essentially reconstituting miR-187 levels when they are pathologically decreased in sepsis.

The association between miR-187 levels and human sepsis as well as sepsis-induced cardiac dysfunction was evaluated in 3 separate patient cohorts. In human studies we showed that miR-187 is contained and its abundance (copy number) decreased in circulating exosomes from septic patients. We were also able to measure and visualize the decrease in miR-187 in the LV from patients who died with sepsis compared to patients who died from other causes unrelated to sepsis or cardiac disease (e.g. cancer). In septic patients with echocardiographic documented heart function, low miR-187 levels were associated with a higher sequential organ failure assessment score and the neutrophil to leukocyte ratio. Importantly, in septic patients with normal hearts, low miR-187 levels were associated with a high lactate and lower mean arterial pressure suggesting this may herald future worsening cardiac function. The limitation of these studies is that these were observational cohorts, therefore, potential bias was not randomly distributed. Moreover, while we only looked at patients with echocardiogram proven abnormal cardiac function, and we excluded patients with a previous history of cardiac disease, we could not completely exclude the possibility that some patients had a priori undiagnosed cardiac dysfunction. Our results do raise the possibility that, in the future, circulating miR-187 levels may represent an effective strategy to select patients for therapy.

The role of miR-187-3p is still under investigation, with limited studies focusing primarily on its overexpression in cancer cells.[30–32] Only one study had previously suggested miR-187’s potential to reduce TNFα production in monocytes.[33] Previous data from our group identified miR-187 as a key miRNA with potential therapeutic implications for sepsis-induced cardiac dysfunction and the beneficial effects of MSC administration in increasing miR-187 levels.[7] Our initial investigations using MSCs for the treatment of sepsis, revealed differential miRNA expression in the septic myocardium, positioning miR-187 as a promising candidate for further study. The conservation of miR-187 across all mammals underscores its evolutionary importance, suggesting a fundamental role in the regulation of critical immune and physiological responses.[34] Of translational relevance, we also found a possible trend for increased miR-187 levels in the 9 septic patients that received MSCs in the first clinical trial of MSCs for the treatment of severe sepsis.[59] While the sample size was small, the trends observed aligned with the patterns seen in our CLP mouse model.

Mechanistic studies strongly support an anti-inflammatory role for miR-187 that is pathologically suppressed during sepsis. miR-187 levels in both human and mice are usually elevated at baseline and plumet early in the course of sepsis. Decreased circulating and LV tissue miR-187 levels may liberate proinflammatory mediators TNFα and IL-6 from miR-187-induced post-transcriptional inhibition. We also suspect that miR-187 may directly or indirectly reduce the expression and potential secretion of (cardiac) alarmins S100A1, S100A8, S100A9, S100A4, thus attenuating DAMP activation and its contribution to sepsis-induced organ failure. A limitation of this study is that we did not fully explore this hypothesis. This line of inquiry is the focus of future studies. Here, we focused on showing that delivery of miR-187 to the heart reduces expression levels of transcription factors MyoD, TBX15 and Mef2c preventing sepsis induced heart failure remodelling to support the hypothesis that the heart, and cardiomyocytes, are active participants in the pathophysiology of sepsis and just not innocent by-standers.

A major limitation to drug development for sepsis is the enormous complexity (quantity) and heterogeneity (quality) of the variability. Hundreds, if not thousands, of genes, proteins and cells are altered during sepsis, and identifying changes that are causatively linked to organ dysfunction and mortality is challenging. We used our knowledge from regenerative medicine to select a candidate therapeutic miR that - based on a series of in vitro and in vivo experiments - we postulated would reduce LV dysfunction and improve mortality. We demonstrate that IV MSC administration in a preclinical model of sepsis reduces inflammation, improves myocardial dysfunction and reduces overall mortality. We took EVs derived from clonally expanded and immortalized MSCs and compared their miR content and effect to control platelet derived EVs and showed that these EV particles derived from MSCs were able to recapitulate the anti-inflammatory effect and prevent LV dysfunction and overall mortality from sepsis. Sequencing of EV-miRs identified miR-187 as a critical anti-inflammatory cardiac miR. In the process of investigating miR-187, we also found that IL-6 is a novel target of this miR, establishing miR-187 as a critical inhibitor of IL-6 mediated contractile dysfunction in sepsis. [35–37]

Based our data, we postulated that delivery of miR-187 could attenuate inflammatory cytokine production, enhance cardiac function, and improve overall survival. However, our studies do not prove that miR-187 contained in MSC-derived EVs are the actual “active ingredient” explaining the therapeutic paracrine effects of MSC-derived EVs. Whether microRNAs contained in EVs have any therapeutic relevance at all is a controversial and active area of research.[38] From our perspective, our studies are not intended to prove that miR-187 detected in our MSC-derived EVs are the active ingredient, this was simply the approach we used to identify miR-187 as a possible novel therapeutic target. To indirectly evaluate whether EV-miR-187 is in fact therapeutically relevant, we treated septic mice with both MSC-EVs with or without LNP-miR-187 mimic or LNP-miR-187 inhibitor. Co-treatment of MSC-EVs with LNP-miR-187 inhibitor could not abrogate the beneficial effects of MSC-EVs. In parallel, MSC-EVs co-administration with the LNP-miR-187 mimic reduced the therapeutic benefit of LNP-miR-187 alone. While not definitive, these findings suggest that miR-187-3p is an important, but not exclusive, effector of MSC-EV–mediated benefit. That is to say, MSC-EVs contain other properties (nucleic acids, proteins or/and lipids) that both partially prevent the detrimental effects of the miR-187 inhibitor and partially abrogate the beneficial effects of the LNP-miR-187 mimic delivery. This data is intriguing, and sets the stage for future development of designer-engineered EVs containing a cocktail of well selected ingredients for the treatment of treatable traits in sepsis derived from regenerative medicine studies.

In conclusion, our data supports the safety and efficacy of miR-187 delivery as a potential strategy for the treatment of sepsis-induced organ failure with a specific focus on cardiac involvement. Future studies in larger animal, or/and non-human primates should provide supportive evidence for first-in-human clinical studies of safety and efficacy.

## Materials and Methods

### MSCs culture and EVs isolation

A frozen vial of human bone marrow-derived MSCs were graciously donated by Dr. Darwin Prockop, (Texas A&M Health Science Center). Cells were thawed, expanded, and cultured according to the previously published literature. [39, 40] MSCs in all in vivo experiments were used between passages 3-8. The capacity of these MSCs to differentiate into adipocytes, osteocytes, and chondrocytes was previously demonstrated and published [40]

Primary human bone marrow clonally expanded and immortalized MSCs (ciMSCs) were cultured and characterized as previously described [41, 42], according to criteria established by the International Society for Cellular Therapy. [43]

EVs were prepared and characterized according to published minimal information criteria for studies of extracellular vesicles [44]. In total, 2 × 10^7^ cells were seeded in Nunc EasyFill Cell Factory System (ThermoFisher Scientific, Waltham, MA) and raised in 400-ml culture medium. Once MSCs reached approximately 50% confluency, the culture media were removed, and conditioned media (CM) were harvested every 48 h until passaging (80–90% confluency). CM were cleared from residual cells and debris by 2000 × g centrifugation for 15 min (Rotor: JS-5.3; Beckman Coulter, Germany), and supernatants were stored at − 20°C until further processing. CMs were screened regularly for mycoplasma contamination (VenorGeM OneStep, Minerva Biolabs, Germany). EVs were prepared from thawed pooled CMs according to our standard procedure, i.e., by polyethylene glycol 6000 (PEG) precipitation followed by ultracentrifugation [45, 46]. EVs were solved in 10-mM HEPES 0.9% NaCl buffer (ThermoFisher Scientific) and stored at − 80 °C as 1-ml aliquots containing EVs harvested from CMs of 4 × 10^7^ MSCs. The batch of ciMSC-EV 41.5 clone 6 used in this study was prepared from a 1.2-l pool of CMs harvested from passages 34 to 37 corresponding to a total cell equivalence of 1.92 × 10^8^ cells. The primary MSC-EV 41.5 batch was prepared from a 4.4-l pool of passages 4 to 5 CMs corresponding to a cell equivalent of 6.45 × 10^8^ cells. As a negative control, platelet-derived EVs (Plt-EVs) were prepared from human platelet lysate (hPL)-supplemented media processed in parallel and using the same protocol as for ciMSC-EVs, including all incubation, PEG precipitation, centrifugation, and resuspension steps. The specific batch used, hPL-EV 07/20, had a protein concentration of 4.2 µg/µL and a particle concentration of 4.9 × 10⁹ particles per µL. Characterization of classical tetraspanin markers (CD9, CD63, CD81) was not available at the time of this study. The Plt-EVs were prepared and analyzed by Robin Dittrich (TA, Bernd’s group), and further information regarding marker profiling may be available from Bernd.

### Lipid Nanoparticle Production

For LNP formulation, the aliquots were then dried with inert gas (N2 or Ar) then placed in a vacuum chamber overnight to remove residual solvent. Films were then stored at-20°C. On the day of formulation, the lipid film was dissolved in ethanol at a concentration of 6.8 mM (3.6 mg total lipid in 1 mL ethanol). The miRNA solution was prepared by suspending 5 nmol miRNA mimic (miR187a-3p mercury LNA mimic, cat#339174, Gene Globe ID:YM00470613-AGA, Qiagen, Alameda CA) inhibitor (miR187a-3p, cat#339174, Gene Globe ID:YI04109154-AGA, Qiagen) or scrambled control (mimic cat# YM00479902-AGA, inhibitor cat#YI0019906-AGA, Qiagen) in 1200 μL 0.1 M acetate buffer (pH 4). The miRNA and lipid solutions were then mixed via microfluidic mixing on a NanoAssemblr Benchtop (Precision Nanosystems) with a flow rate ratio of 3:1 respectively, and a total flow rate of 9 mL/min. After mixing, the LNPs were diluted to 12 mL with PBS (pH 7.4) and were purified and by filtration with a 30 kDa cut-off centrifugal filter before resuspension in PBS, then sterile-filtered through a 0.45-micron PTFE syringe filter. A 50μL aliquot was collected and diluted 20-fold in PBS and size, polydispersity index (PDI) and zeta-potential were then measured on a Zetasizer Nano ZS (Malvern Panalytical). The concentration and median size of the LNPs were determined by Nanoparticle Tracking Analysis (ZetaView, ParticleMetrix) at a dilution factor of 2×10^5^. The LNPs were then stored in PBS at 4°C prior to use. The protocol used was adapted from published protocols. [47]

Following filtration of LNPs, encapsulation efficiency was measured. Loading of LNPs was quantified using a Quanti-iT RiboGreen assay (Invitrogen) according to manufacturer’s instructions. To ensure accurate quantification, a standard curve was created utilizing the same RNA which was loaded into the LNPs. Fluorescence readings were collected on a CLARIOstar Plus reader (BMG Labtech). Encapsulation efficiency (EE) was calculated using the following formula:

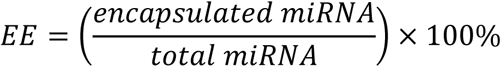

Formulated LNPs had an average concentration of 5-6 x 10^12^ LNPs/mL, with a final encapsulation efficiency of 80-95% (lower range for miRNA inhibitors, and higher for miRNA mimic), with an average size of 110 nm, a PDI of 0.15 and a zeta-potential of-2 to-3 mV.

### Cell viability

THP1 cells were plated at 2.5×10^5^ /mL in RPMI 1612 media containing no serum overnight, then treated with miR-187 LNP at concentrations 0-1 nM for 24 hours. Cell viability was assessed using the Alamar blue assay kit (cat#AO50100, ThermoFisher Inc.) according to the instructions of the manufacturer.

### Cell cytotoxicity

THP1 cells were plated and treated as above. Lactate dehydrogenase (LDH) is a is a well-defined and reliable indicator of cytotoxicity. Cell cytotoxicity was assessed using the CyQUANT™ LDH Cytotoxicity Assay Kit (cat#C20300, ThermoFisher Inc.) according to the instructions of the manufacturer.

### Cell apoptosis

THP1 cells were plated and treated as above. Cell apoptosis was assayed using the eBioscience™ Annexin V Apoptosis Detection Kit (cat#88-8005-72, TherfoFisher Inc.) according to the instructions of the manufacturer. The kit utilizes viability dye-conjugated annexin V to label phosphatidylserine (PS), an early marker of apoptosis, on the extracellular membrane.

### Cryo-Electron Microscopy of Lipid Nanoparticles

LNP-miR-187 samples were prepared on 300 mesh Quantifoil R2,2 grids (EMS). Grids were first charged for 30 seconds with a Pelco EasiGlow (Ted Pella) glow discharge cleaning system, followed by application of 4.0 µL of LNP solution (6 x 1012 LNPs/mL). Grids were blotted and plunge-frozen in liquid ethane using a FEI Vitrobot IV. Vitrobot temperature was set to 4°C, humidity set to 100%, blot time was set to 4.5 seconds, wait time was set to 5 seconds, blot force was set to 2 (arbitrary units), blot total was set to 1, and drain time was set to 0 seconds. Samples were stored in liquid nitrogen until imaging.

Samples were loaded into the TEM using the Gatan 626 single tilt cryo-EM holder. All images were taken on a FEI Talos L120C TEM equipped with BM-Ceta metal-oxide semiconductor camera. Images were taken using 120kV at 57000x magnification corresponding to a pixel size of 249 pm. Images were collected at a defocus of –3.5 to-1.5 microns using the FEI TEM Imaging & Analysis (TIA) software.

### Animal Protocol

All studies were approved by the animal care committee at St. Michael’s Hospital in accordance with Canadian Council of Animal Care guidelines. Both male and female C57BL/6 mice (8–12-week-old), weighing 20–28 g, were purchased from Jackson Laboratories (Bar Harbor, ME) and housed in our facility with standard 12-hour light/dark cycle with *ad libitum* access to food and water. Only for the MSC study, we used exclusively male mice. Mice were acclimatized to the animal facility for at least a week before beginning experiments. Experiments were conducted in accordance with minimum quality threshold in pre-clinical sepsis studies. [48]

### Cecum ligation and puncture (CLP) model

Mice were randomized to sham or cecal-ligation and puncture (CLP) procedure as previously described (Blue journal paper). Briefly, mice were anesthetized with isoflurane anesthesia (1.5– 4 vol%, total duration of surgery: 5-7min). A midline laparotomy was performed the cecum was exposed, ligated with 3-0 cotton thread below the ileocecal valve, and punctured four times with a 21-gauge needle. In the sham group, an abdominal incision was made, the cecum was exposed, mobilized, but there was no cecal ligation or perforation. Both layers of the abdominal cavity were closed. The animals were returned to their cages, with free access to water and food, given subcutaneous fluid resuscitation (saline equivalent to 30 ml/Kg every 12 h), buprenorphine (0.1 mg/kg every 48 h) and Imipenem (25 mg/kg, Ranbaxy, Pharmaceuticals Canada Inc.), intraperitoneal (IP) 6h after surgery and then once a day.

*Interventional Studies*: Six hours after surgery, CLP and sham mice were further randomized to receive sterile saline (negative control) vs treatment or placebo, as appropriate (detailed for each interventional study below). All treatments were single dose, delivered intravenously (IV) via the retro-orbital vein (RO), except the MSC injection which occurred via the tail vein. RO injection is minimally invasive and avoids first-pass metabolism in the liver – like IV infusion through the jugular vein using a central venous line in humans. All assessors were blinded to treatment group assignment. Unless otherwise specified, all treatments (cells, EVs and LNPs) were diluted in saline and delivered in equal volume (100 μL).

A total of 8 separate experiments, evaluating three separate and independent treatments were conducted:

i. MSC administration: Groups were sham, CLP+sterile saline, and CLP+MSCs (2.5×10^5^ cells). Dose of MSCs was as previously published. [49]
ii. MSC-derived EV administration: groups were sham, sham+ciMSC-EVs, sham Plts-EVs, CLP+sterile saline, CLP+ciMSC-EVs, CLP+Plts-EVs (5×10^4^ cell equivalents per gram of bodyweight). The dose for both types of EVs was as previously described [41, 45, 50].
iii. LNP-miR-administration: groups were sham, CLP+sterile saline, CLP+scrambled (SCR) miR as a negative control, and CLP+miR-187 mimic LNP, CLP+miR-187 inhibitor, CLP+empty LNPs. Doses were 1 nmol per mouse. Dose of miR and scrambled were based on previously published work using liposomal delivery.[11, 12]

*Monitoring and welfare assessments*: Mice wellbeing was monitored every 4h. Cage enrichment accessories, such as a mouse swing and cottage, were used to help mitigate anxiety and stress reactivity. Welfare score was calculated based on a seven-point scale (total of 28) evaluating gross appearance, hydration status, motor activity and reflexes as well as respiratory rate and quality, body weight, rectal temperature, and glucose levels. Mice were humanely euthanized if at any given time point the welfare score was greater than 21, or if the points ascribed to respiratory parameters were more than 3. Results were reported as probability of survival. Assessment of physiological parameters were performed by an experimenter blinded to treatment assignment, and welfare assessment of the mice was performed in accordance with the St. Michael’s Hospital Vivarium Facility. All mice were randomized to experimental groups by providing a unique ID number per mouse in experiments, and were assigned to treatment groups using a random number generator (https://www.calculatorsoup.com) in accordance to the Animal Research Reporting of In Vivo Experiments guidelines.[12]

*End-points:* Animals were sacrificed by anesthetic overdose (isoflurane) when they met criteria for euthanasia or at the endpoint of each study.

### Echocardiography

Transthoracic echocardiography was performed in mice anesthetized with 1-2% isoflurane at 40 or 72 hrs after the CLP procedure. Two-dimensional echocardiography (Hewlett-Packard Sonos 5500, Philips Ultrasound, Bothel, WA) was performed using a broadband 5-to 12-MHz ultrasound probe (S12 transducer). Animals were imaged in the supine position. A standoff was created by using a small water bath. The imaging depth was set at 2 cm and magnified to 1 cm, allowing a frame rate of 100 –120 Hz. A short-axis two-dimensional view of the left ventricle (LV) at the midpapillary muscle level was digitally acquired and stored for off-line analysis. End diastole was defined as the frame with the largest cavity size, and end systole was defined as the frame with the smallest cavity size. LV internal diameters at end diastole (LVIDd) and end systole (LVIDs) were measured on the two-dimensional image using the standard leading edge to leading edge technique. Each measure was made by a single observer blinded to treatment assignment. For each measurement, three consecutive cardiac cycles were measured and then averaged. Per cent fractional shortening (FS) was calculated as (LVIDd - LVIDs)/LVIDd x 100, and per cent ejection fraction (EF) was calculated as (LVIDd^3^ - LVIDs^3^)/LVIDd^3^ x 100.[51]

### Hemodynamic measurements

In mice maintained under 2% isoflurane, systemic systolic and diastolic arterial pressures were measured using a Millar micro-tipped catheter transducer (modelSPR-671, Millar Instruments, Houston, TX) inserted into the right carotid artery. The catheter was then advanced into the LV for the measurement of LV pressure, LV end-diastolic pressure, and the maximum rate of pressure rise (+dP/dtmax) and decline (-dP/dtmax). Pressure signals were recorded using a Power Lab data-acquisition system and analyzed via Chart 5 software (AD Instruments, Colorado Springs, CO).[51]

### Histology

The right lung, right kidney, liver, and a central full-thickness slice of the wall of the heart obtained by 2 transverse cuts at levels corresponding to the one-third and two-thirds of the length of the heart, were removed, fixed in 10% neutral-buffered formalin, embedded in paraffin, cut (4 μm thick) longitudinally from the central zone with a microtome and stained with hematoxylin-eosin (H&E) for histologic analysis. Photomicrographs were obtained using a Zeiss AxioScan 7 SlideScanner in bright field mode (Zeiss, Germany) at 200x magnification. Lungs were assessed using morphometric standards established by American Thoracic Society and European Thoracic Society [52]. Three random regions from each slide were evaluated at a magnification of ×200 across 10 random, noncoincident microscopic fields. In the heart, lymphocyte infiltration was assessed by counting the lymphocytes between the cardiomyocytes using the Weibel method [53]. Three random regions from each slide were evaluated. Lymphocytic infiltrate between cardiomyocyte fibers with discreate cardiomyocytes necrosis.

To assess the biodistribution and retention of lipid nanoparticles (LNPs), in vivo fluorescence imaging was performed using a Newton 7.0 Imager (Vilber Lourmat Deutschland GmbH). Mice were injected intravenously with Cy5-labeled LNPs at 6 hours post–cecal ligation and puncture (CLP) or sham surgery. Organ-specific and Whole-body images were acquired at multiple time points (24, 48, 72 hours and Days 1–14 post-injection respectively) under isoflurane anesthesia. Fluorescence was detected in the Cy5 channel with a fixed exposure time of 10 milliseconds. Images were captured using the device’s standard acquisition settings and analyzed with Bio-Vision+ software (Vilber). Fluorescence intensity was quantified in specific regions of interest (ROIs) including the heart, liver, lungs, spleen, and kidneys, and presented as arbitrary units (a.u.).

### Electron microscopy

Samples were initially fixed with 4% Paraformaldehyde plus 1% glutaraldehyde in 0.1M phosphate buffer (pH 7.2) for 2 hours at room temperature and then post-fixed with 1% osmium tetroxide in 0.1M phosphate buffer for 1 hour at room temperature Then, dehydration was performed using a graded series of ethanol. Samples were permeated with EPON resin and placed in a BEEM embedding capsule and polymerized at 60°C for 48 hours.

After complete polymerization, the solid resin block containing the sample was sectioned on a Reichert Ultracut E microtome to 90nm thickness and collected on 200 mesh copper grids. Sections were stained using saturated uranyl acetate and Reynold’s lead citrate. The stained sections were examined and photographed on a FEI Talos L120C transmission electron microscope. Mitochondria were counted using the Weibel method.[53] The Z line distance and mitochondrial area were measured using Image Pro Plus 7 software (Media cybernetics, Rockville, MD).

### Cardiomyocyte Isolation and Transfection

Neonatal cardiac myocytes were isolated from the ventricles of 2-day-old control C57BL/6 mice [22]. Cells in DMEM F-12 medium (Thermo Fisher Inc.) supplemented with 10% FBS and 5% Pen/Strep (ThermoFisher, Inc.) were cultured at a concentration of 5×10^5^ in 60 mm plates for 24 hrs at 37°C. Media was replaced with antibiotics-free media 12 hours before transfection. Cells were transfected with Hiperfect transfection reagent (ThermoFisher Inc.) complexed with LNA miR-187 mimic (10 nM, Qiagen) or negative control (10 nM, Qiagen) according to the supplier’s instructions for 24 hrs. The media was then removed and replaced with transfection reagent-free media treated with or without lipopolysaccharide (LPS) (1 μg/mL, L5543, Sigma-Aldrich, Burlington, MA) for an additional 24 hours. RNA and protein were isolated as described below.

### Dynosore experiment

Neonatal cardiac myocytes were isolated as described above. Cells were pretreated with Dynosore (10 mM final, D7693, Sigma-Aldrich, Burlington, MA) [54] for 4 hrs. LPS (1 μg/mL) was added together with EV-ciMSCs or EV Plts (2.5 or 5 μL). Cells were harvested 24 hours after EVs addition. RNA and protein were isolated as described below.

### RNA and microRNA isolation

RNA was extracted as previously described.[7] Total RNA was isolate from whole heart tissue or cells using Trizol (ThermoFisher Inc.) and purity and integrity of RNA were checked by the 260/ 280 nm absorbance ratio using NanoDropTM (ThermoFisher Inc.). Only preparations with ratios higher than 1.8 and no signs of RNA degradation were utilized. RNA concentrations were determined by absorption at 260 nm.

### Polymerase chain reaction

For cDNA synthesis, 1 μg of total RNA was used with the SuperScript IV First-Strand Synthesis System (ThermoFisher Inc.). Expression analyses were performed by qRT-PCR using Power SYBR Green Master Mix (ThermoFisher Inc.) on a QuantStudio 7 Flex Real-Time PCR System (ThermoFisher Inc.). The average threshold cycle (Ct) value from three technical replicates was used for each sample. RNA18S and U6 served as loading controls for mRNA and miRNA, respectively. Data were analyzed using the 2^ΔCt method described by Schmittgen and Livak, 2008 [55], and relative mRNA data are presented as the mean ± standard error of the mean (SEM).

Gene specific sequences of oligonucleotide primers (proprietary sequences - IL-1b, IL-6, TNFa, S100A8, S100A9, IL-10, S100A1, NppA, Myh1, Myh6, Myh7, Myh11, Casq2, Titin, Mef2c, TBX15, MyoD1, 18S, GAPDH, miR-187-3p, U6 snRNA; Qiagen, Alameda, CA) based on DNA sequences in the National Center for Biotechnology database were purchased from Qiagen. Real time quantitative RT-PCR was performed as per manufacturer’s protocol (Qiagen). Briefly, a 25 mL reaction volume containing 12.5 mL of RT^2^ SYBR Green qPCR Master Mix, 10.5 mL of H_2_0, 1 ml of gene-specific 10 mM qRT-PCR primer pair stock of gene of interest.

### Blood chemistry analysis

Whole blood was collected from mice and allowed to clot at room temperature. Serum was isolated by centrifugation and submitted to The Centre for Phenogenomics (Toronto, Canada) for comprehensive clinical chemistry analysis, including measurement of up to 18 analytes using their murine panel.

### ELISA

Plasma was isolated from whole blood and used for the determination of murine/human TNF-a, human IL-1b (DuoSet ELISA kit, R&D Systems Inc., Minneapolis, MN), murine IL-6 (ProteinTech, Rosemont, IL), and cardiac troponin (Cusabio, Houston, TX) were quantified according to the manufacturer’s instructions. The sensitivity for each kit was as follows: TNFα - 31.2 pg/mL, IL-1β - 3.9 pg/mL, IL-6 - 3.8 pg/mL), cardiac troponin - 3.9 pg/mL).

### 3’UTR – Luciferase assay

All luciferase reporter constructs were purchased from Switch-Gear Genomics (Menlo Park, CA). The 3’UTR reporter construct of IL-6 gene (cat#S803848) comprised the 3’UTR of the human IL-6 gene cloned downstream of a constitutive ribosomal protein L10 promoter and the Renilla luciferase gene reporter. Neonatal murine cardiomyocytes were cultured in a 96 well plate (as described above) and co-transfected with the SwitchGear GoClone reporter IL-6 luciferase construct in combination with either miR-187 mimic or scrambled miR and incubated for 24 hrs. Cells were then treated with LPS 1 μg/mL or vehicle (PBS) for an additional 24 hrs. LightSwitch Luciferase Assay Reagent (Switch-Gear Genomics, cat# LS010) was added to each sample and luciferase activity was measured three times using the BioTek Synergy/neo microplate reader (Agilent, Santa Clara, CA). All results were normalized against a blank control.

### Cardiomyocytes mRNA sequencing

Cardiomyocytes were isolated from 1–3-day-old neonatal mice and transfected with 50 nM miR-187-3p mimic or miRNA negative control using HiPerFect transfection reagent. After 24 hours, cells were exposed to either LPS (1 µg/mL) or fresh media for an additional 24 hours. Total RNA was extracted, and miRNA integrity was assessed using an Agilent Bioanalyzer. High-quality samples were submitted to **The Centre for Applied Genomics (TCAG, Toronto, Canada)** for bulk mRNA sequencing.

### EVs miRNA sequencing

ciMSC-and plts-derived EVs were generously provided by the Giebel laboratory (University of Duisburg-Essen, Germany). These samples were sent to HTG Laboratory Services (Houston, Arizona) for RNA isolation and miRNA sequencing using the HTG EdgeSeq miR Whole-Transcriptome Assay. The raw miRNA count data obtained were subjected to differential expression analysis to identify miRNAs showing differences between the two EV groups. All statistical procedures were performed using R software (version 4.2.2), and Bioconductor (edgeR and DESeq2) was used for analysis of differential gene expression.[56, 57]

### Ethics approval – Human Samples

All patients were enrolled under local ethical board approval. Informed written consent was obtained upon enrollment from the patient or their legal representative.

### FFPE human samples

Archival blocks of formalin fixed paraffin embedded (FFPE) heart tissues from N=6 patients who died with sepsis (3 males and3 females; mean age 54.8 years) and 6 control patients deceased from cancer (non-septic hearts: 3 males and 3 female; mean age 57.3 years) were obtained through the 1^st^ Department of Pathology, University of Athens Medical School, Athens, Greece (REB#10-2021). RNA was extracted from FFPE human sections using the RNeasy FFPE kit (QIAGEN, cat. no. 73504) according to the manufacturer’s instructions.

### miRScope

Quantification of miR-187a-3p in heart tissue was performed using miRNAscope in situ hybridization with the miRNAscope HD Detection Reagent-RED (Cat. No. 324510, Advanced Cell Diagnostics). Positive controls included the SR-RNU6-S1 control probe (Cat. No. 727871-S1) and the SR-Mm-Snord-85-S1 probe (Cat. No. 728951-S1), while the SR-Scramble-S1 probe (Cat. No. 727881-S1) served as a negative control. The miR-187a-3p-specific probe (Cat. No. 1183471-S1) was used for target detection in the miRNAscope HD RED assay. Images were acquired at 63× oil immersion (1.4) using a ZEISS wide-field microscope with deconvolution.

### INSIST cohort and healthy volunteers

Septic patients admitted to the Medical Surgical Intensive Care Unit (MSICU) of St. Michael’s Hospital, Toronto meeting the clinical criteria for sepsis 3 and diagnosed with a Multiple Organ Dysfunction Score (MODS) were enrolled into the Cellular and Molecular Mechanisms of Prolonged Neutrophil-Mediated Inflammation in Trauma and Sepsis (INSIST) study (REB#09-216, including healthy volunteers). Inclusion criteria for patient enrollment - ICU admission ≤ 48 hours, suspected infection + 3 positive SIRS criteria in the past 24 hours (temperature > 38 C or < 36 C, heart rate > 90 BPM, PaCO2 < 32, WBC >12, MODS 4 55 56). Exclusion criteria - Corticosteroids (> 5 mg/day prednisone or equivalent) or biologic response modifier within past 7 days, transfusion > 2 units in past 24 hours, lack of commitment to full ICU support (excluding CPR), and lack of informed consent.

Whole blood was collected in heparinized tubes from patients with sepsis (n=14) and healthy volunteers (n=6). Red blood cells were allowed to separate from whole blood for 45 minutes at room temperature in the dark. The plasma and white cell content was collected using a sterile serological pipette and placed on Ficoll and centrifuged at 400 g for 20 minutes at 22 °C. Plasma fractions were collected in 1.5 mL Eppendorf tubes and stored at-80 °C.

### Exosome isolation

We isolated exosomes from 1 mL of healthy and septic patient blood using the ExoEasy kit according to the manufacturer’s instructions (Qiagen). The isolated exosomes at a volume of 200 μL were characterized by Nanosight technology, electron microscopy, and qRT-PCR for GT130 (negative), and CD9, CD63, CD81 (positive) exosome markers as previously described.[58] Exosome-derived miRNA was isolated using Qiagen miRNeasy Kit according to the manufacturer’s protocol as described. Real time quantitative RT-PCR was performed as per manufacturer’s protocol (Qiagen). Briefly, a 25 mL reaction volume containing 12.5 μL of miRCURY LNA SYBR Green Master Mix, 10.5 mL of H_2_0, 1 μl of miR-specific 10 mM qRT-PCR primer pair stock of gene of interest.

### CISS Cohort and healthy volunteers

The Cellular Immunotherapy for Septic Shock (CISS) trial was approved by Health Canada and the Ottawa Health Sciences Network Research Ethics Board (OHSN-REB No.: 20140809-01H) and registered on clinicaltrials.gov (NCT02421484). Details on trial design, participant recruitment, MSC source and preparation, and trial outcomes have been described previously.[59] Briefly, the observational cohort consisted of 21 septic shock participants who met CISS eligibility criteria but did not receive MSCs, and the interventional cohort (n = 9) comprised three cohorts of three septic shock participants that received 0.3 (low dose), 1.0 (mid dose), or 3.0 (high dose) million freshly cultured cells per kg body weight, to a maximum of 300 million cells. Peripheral blood samples from healthy participants were obtained with informed written consent at a single center (OHSN-REB No.:2011470-01H).

### COLOBILI cohort and healthy volunteers

Patients recruited and enrolled to the Covid-19 Longitudinal Biomarker in Lung Injury (COLOBILI) trial at Unity Health Toronto were included in accordance with protocol approved by the St. Michael’s Hospital ethics board (REB#: 20-078). Inclusion criteria included patients admitted to the ICU with respiratory deterioration suspected or confirmed to be due to SARS-CoV-2. Only patients found to be COVID19 positive will stay in this cohort. Biological controls included ICU patients found to be COVID19 negative, patients with acute respiratory failure unrelated to COVID-19, and patients admitted to the floor with COVID-19 or acute respiratory failure. Healthy Volunteers recruited healthy volunteers among healthcare workers and allied personnel in the hospital. Exclusion criteria included refusal to participate, inability to record the primary outcomes, failure to obtain blood sample, known to have had COVID in the past. Patients’ data were extracted from the in-hospital electronic medical records, de-identified, and assigned random identification numbers which were used throughout the project.

### Pax Gene tube blood collection

Patient samples were collected in Pax Gene tubes (cat#762165, BD Biosciences, Mississauga, ON) and total RNA was isolated using standard protocol for Qiagen RNAeasy mini kit which allows for isolation of small RNAs.

### Digital droplet PCR

RNA extracted from homogenized whole tissue or cultured cells was reverse transcribed using RT-specific TaqMan primers (cat#4427975, ThermoFisher Scientific Inc.) in conjunction with the TaqMan MicroRNA Reverse Transcription Kit (cat#4366596, ThermoFisher Scientific Inc.). The resulting cDNA was amplified using ddPCR Supermix for Probes (cat#186-3010, Bio-Rad, Hercules, CA) according to the manufacturer’s protocol. Briefly, a final reaction volume of 20 μL, containing 1×master mix, primers and TaqMan probes (cat#4427975, ThermoFisher Scientific Inc.), and 2 μL of cDNA, was loaded onto disposable droplet generator cartridges (before 12.05.2014 cat#186-3008, from 12.05.2014 cat#186-4008, gaskets cat#186-3009, Bio-Rad). A total of 70 μL of droplet generation oil (cat. no. 186-3005, Bio-Rad) was then added, and droplets were formed using the QX100 system (Bio-Rad). The droplets were transferred to a 96-well PCR plate (cat#0030128.613, TwinTec, Eppendorf, Hamburg, Germany), creating ∼20,000 nL-sized water-oil emulsion droplets per sample. The plate was thermocycled to endpoint, sealed, and subsequently placed into the QX100 droplet reader (cat#186-3004, Bio-Rad), following the manufacturer’s recommendations. Fluorescence amplitudes were recorded for all droplets in each sample well using a droplet flow cytometer.

### Statistics analysis

All studies were conducted using a randomized and blinded approach to minimize bias. Observers responsible for assessing endpoints were blinded to treatment assignments. Formal sample size calculations were based on predefined primary outcomes, including mortality rates, levels of inflammatory mediator gene expression, and degree of cardiac dysfunction. Prior to analysis, data were tested for normality using the Shapiro-Wilk test and visual inspection of histograms to determine whether parametric or non-parametric statistical methods were appropriate. Normally distributed variables are expressed as mean and standard error of the mean, whereas non-normally distributed variables are presented as median and inter-quartile range values (IQR; 25th–75th percentile). For two group comparison, the Student’s t-test was used for parametric, and the Wilcoxon Signed Rank Test for nonparametric data. For multiple comparisons, the Kruskal-Wallis one-way analysis of variance (ANOVA) was used for nonparametric followed by the post-hoc Dunn’s test, and the standard one-way ANOVA was applied for parametric data followed by the post-hoc Bonferroni t-test for pair-wise comparisons.

## List of Abbreviations

ALI: Acute Lung Injury
ALP: alkaline phosphatase
ALT: alanine aminotransferase
AST: aspartate aminotransferase
BUN: Blood Urea Nitrogen
Casq2: calsequestrin 2
cDNA: complementary DNA
ciMSCs: clonally-expanded immortalized Mesenchymal Stromal Cells
CISS: Cellular immunotherapy for severe sepsis
CLP: Cecum ligation and puncture
CM: Conditioned media
DAMP: Damage Associated Molecular Pattern
ddPCR: digital droplet Polymerase Chain Reaction
DSPPC: distearoylphosphatidylcholine
EE: Encapsulation efficiency
EF: Ejection Fraction
EVs: Extracellular Vesicles
FDR: False Discovery Rate
FFPE: formalin fixed paraffin embedded
FS: Fractional Shortening
H&E: hematoxylin-eosin
HDL: high-density lipoprotein
IL-10: Interleukin 10
IL-1β: Interleukin 1 beta
IL-6: Interleukin 6
INH: inhibitor
IP: intraperitoneal
IV: intravenous
LDH: lactate dehydrogenase
LDL: low-density lipoprotein
LNP: Lipid nanoparticle
LPS: lipopolysaccharide
LV: Left Ventricle
LVDP: LV end-Diastolic Pressure
LVEF: Left Ventricular Ejection Fraction
LVFS: Left Ventricular Fractional Shortening
LVIDd: Left Ventricle internal diameters at the end of diastole
LVIDs: Left Ventricle internal diameters at the end of systole
Mef2c: Myocyte Enhancer Factor 2C
MIM: mimic
miR/miRNA: microRNA
miR-187-3p: microRNA-187a-3p
Mito: mitochondria
MODS: Multiple Organ Dysfunction Score
MSC-EVs: MSC-derived EVs
MSCs: Mesenchymal Stromal Cells
MSICU: Medical Surgical Intensive Care Unit
Myh1: myosin heavy chain 1
Myh11: myosin heavy chain 11
MyoD: Myogenic Differentiation 1
NFκβ: Nuclear Factor Kappa Beta
NPPA: Natriuretic Peptide Precursor A
PDI: polydispersity index
PEG: polyethylene glycol
Plt-EVs: Platelet derived Extracellular Vesicles
PMA: Phorbol 12-myristate 13-acetate
RIN: RNA Integrity Number
S100A1: S100 Calcium Binding Protein A1
S100A4: S100 Calcium Binding Protein A4
S100A8: S100 Calcium Binding Protein A8
S100A9: S100 Calcium Binding Protein A9
S/C: subcutaneous
SCR: Scrambled
TNFα: Tumor necrosis factor alpha
TXB15: T-Box Transcription Factor 15
UTR: untranslated region
αMhc/Myh6: alpha Myosin Heavy Chain
βMhc/Myh7: beta Myosin Heavy Chain

## Supplemental Figures

**Supplemental Figure 1.**
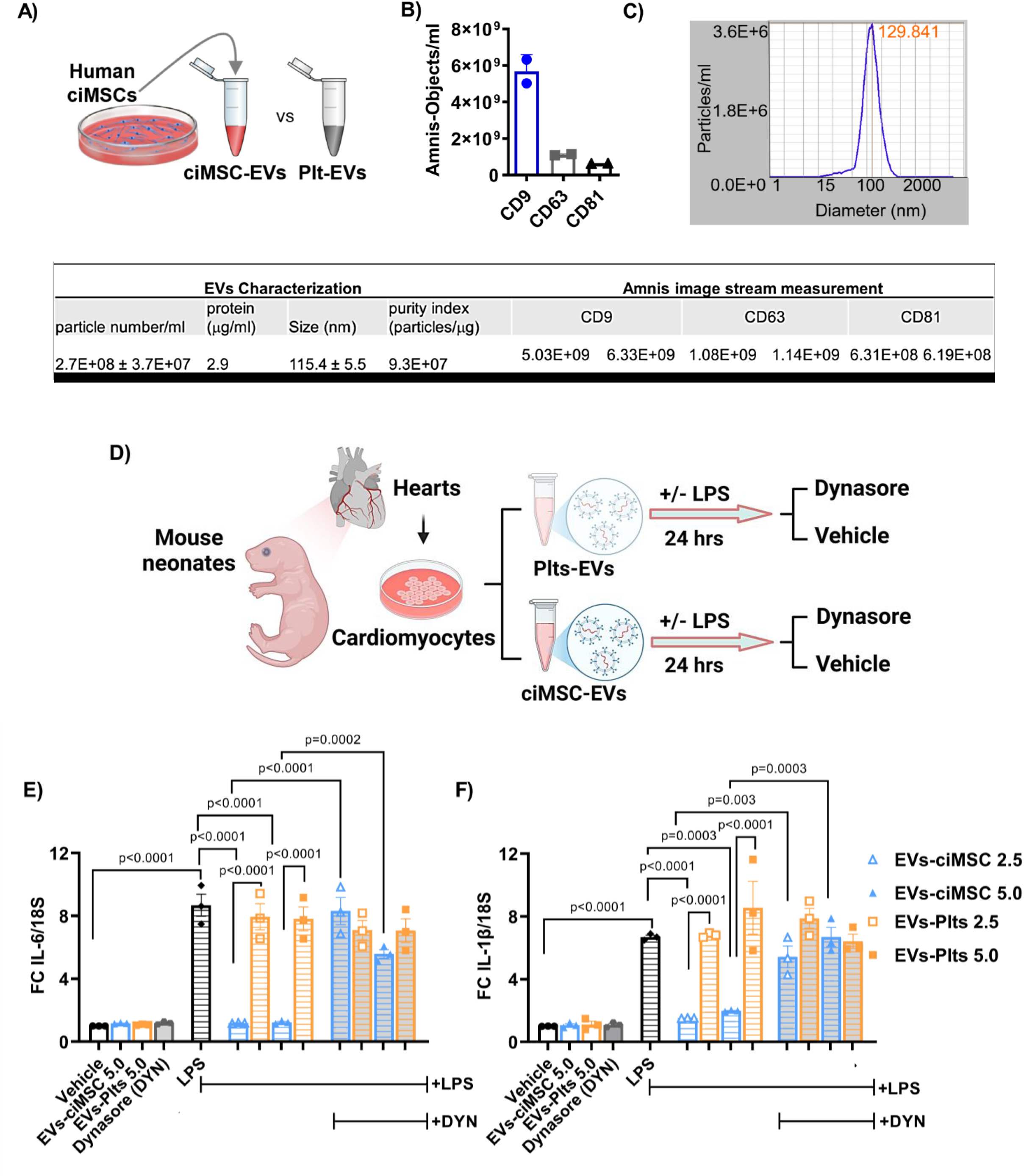
Human ciMSC-EVs attenuate LPS-induced inflammatory signaling in neonatal cardiomyocytes via dynamin-dependent endocytosis. Schematic of extracellular vesicle (EV) isolation from cultured human cardiac-induced mesenchymal stem cells (ciMSCs) or platelet-rich plasma (Plt) (A). Nanoparticle tracking analysis showing expression levels of canonical EV surface markers CD9, CD63, and CD81 (B), and particle size distribution with average diameter (114.4 ± 5.5 nm) (C). Table details of characterization. Experimental workflow showing treatment of primary neonatal mouse cardiomyocytes with ciMSC-EVs or Plt-EVs (2.5 or 5.0×10⁸ particles/mL) in the presence or absence of lipopolysaccharide (LPS, 10 ng/mL) and dynamin inhibitor Dynasore (DYN) (50 µM) for 24 hours (D). Fold change in gene expression of IL-6 (E) and IL-1β (F) normalized to 18S, measured by qRT-PCR. Data are presented as mean±SEM. One-way ANOVA followed by Bonferroni-Dunn post hoc test was used. n=3 per group.

**Supplemental Figure 2.**
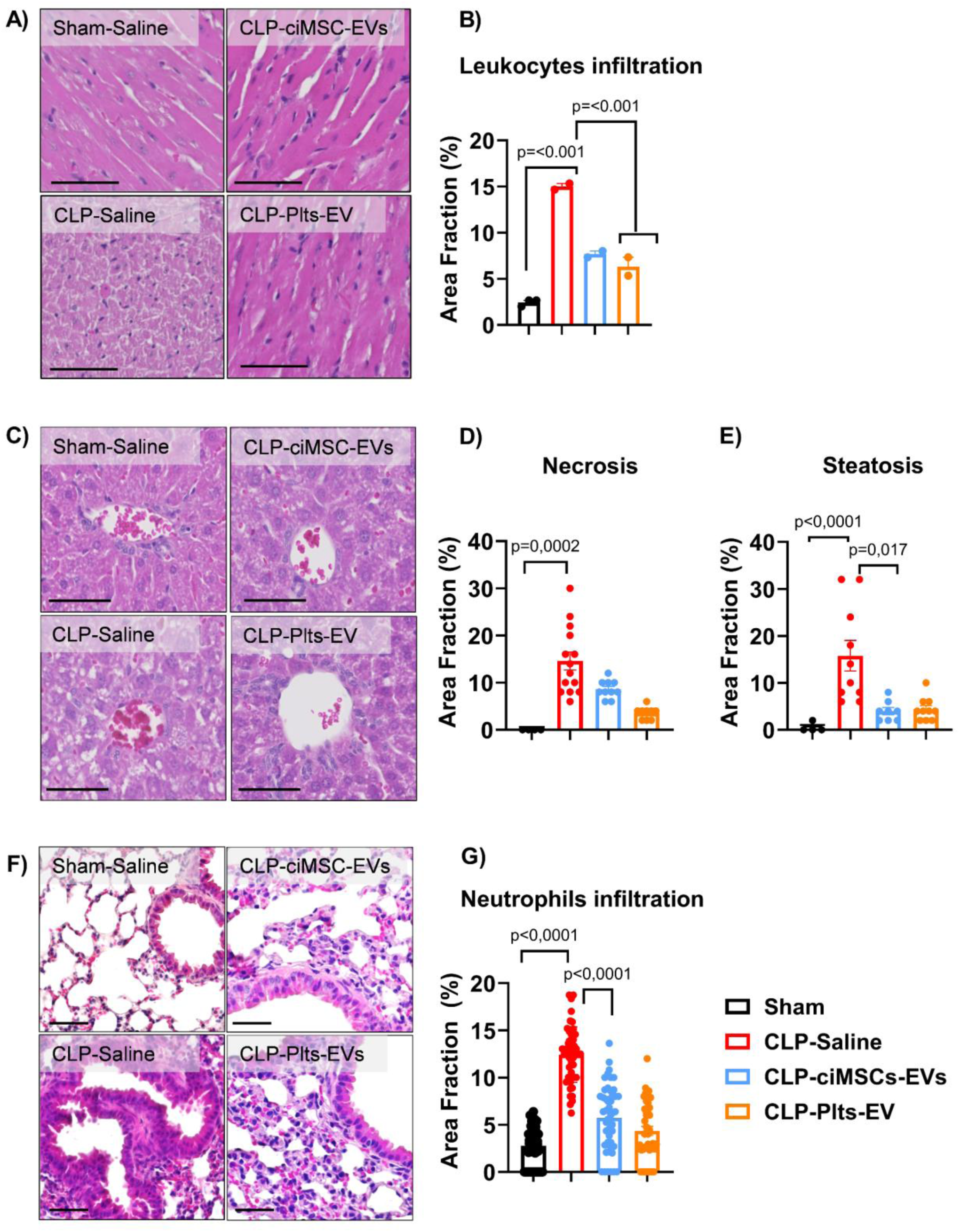
Human ciMSC-EVs reduce leukocyte infiltration, hepatic injury, and lung neutrophil infiltration in septic mice. Representative hematoxylin and eosin (H&E)-stained cardiac sections showing leukocyte infiltration of sham + saline, cecal ligation and puncture (CLP) + saline, CLP + clonally expanded and immortalized mesenchymal stem cell (ciMSCs) extracellular vesicles (EVs), and CLP + platelets (Plts) EVs 48 hrs post-CLP (A), with quantification of leukocyte infiltration represented as mean±SEM of % area fraction affected (B). Representative H&E-stained liver sections from groups as above showing necrotic foci and micro vesicular steatosis (C), with quantification of necrosis (D) and steatosis (E) represented as mean±SEM. Representative lung sections from groups as above stained with H&E showing neutrophil accumulation in alveolar spaces (F), with quantitation of the mean±SEM of % area fraction affected (G). Statistical comparisons performed using one-way ANOVA with Bonferroni-Dunn post hoc test. n=4–10/group.

**Supplemental Figure 3.**
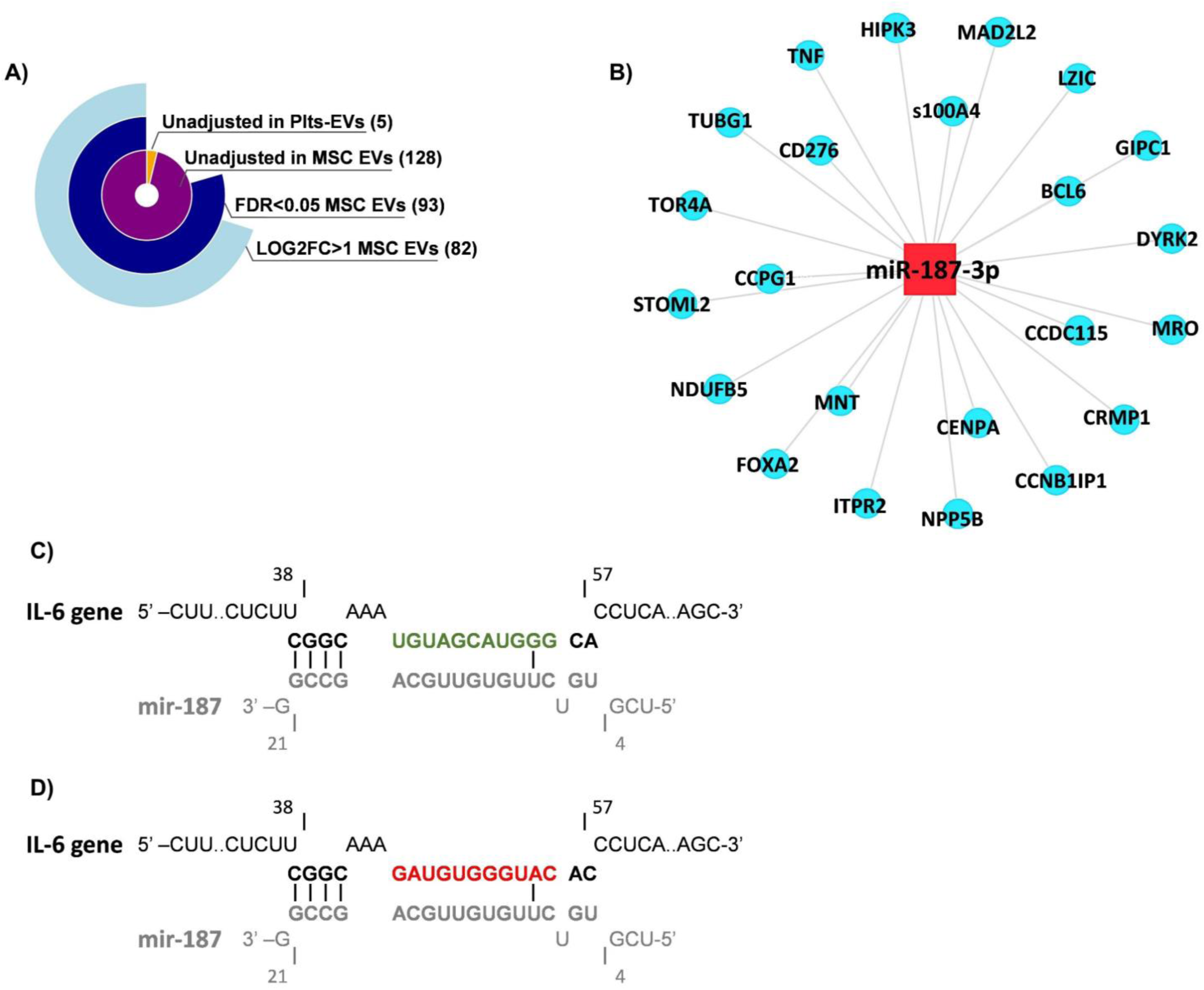
miR-187 selection, known binding targets and seed sequence in 3’UTR of IL-6 mRNA. Circle-pie chart showing process of identification of miR-187 as over-represented in extracellular vesicles (EVs) derived from clonally expanded and immortalized mesenchymal stem cell (ciMSCs) with an adjusted p=value <0.05 and fold change greater that 2 (A). A schematic of miR-187 experimentally supported targets as identified by DIANA-TarBase v9 (B). Binding site sequences between miR-187 (C), scrambled miR (D) and the 3’-UTR of human IL-6 were mapped out using INTARNA.

**Supplemental Figure 4:**
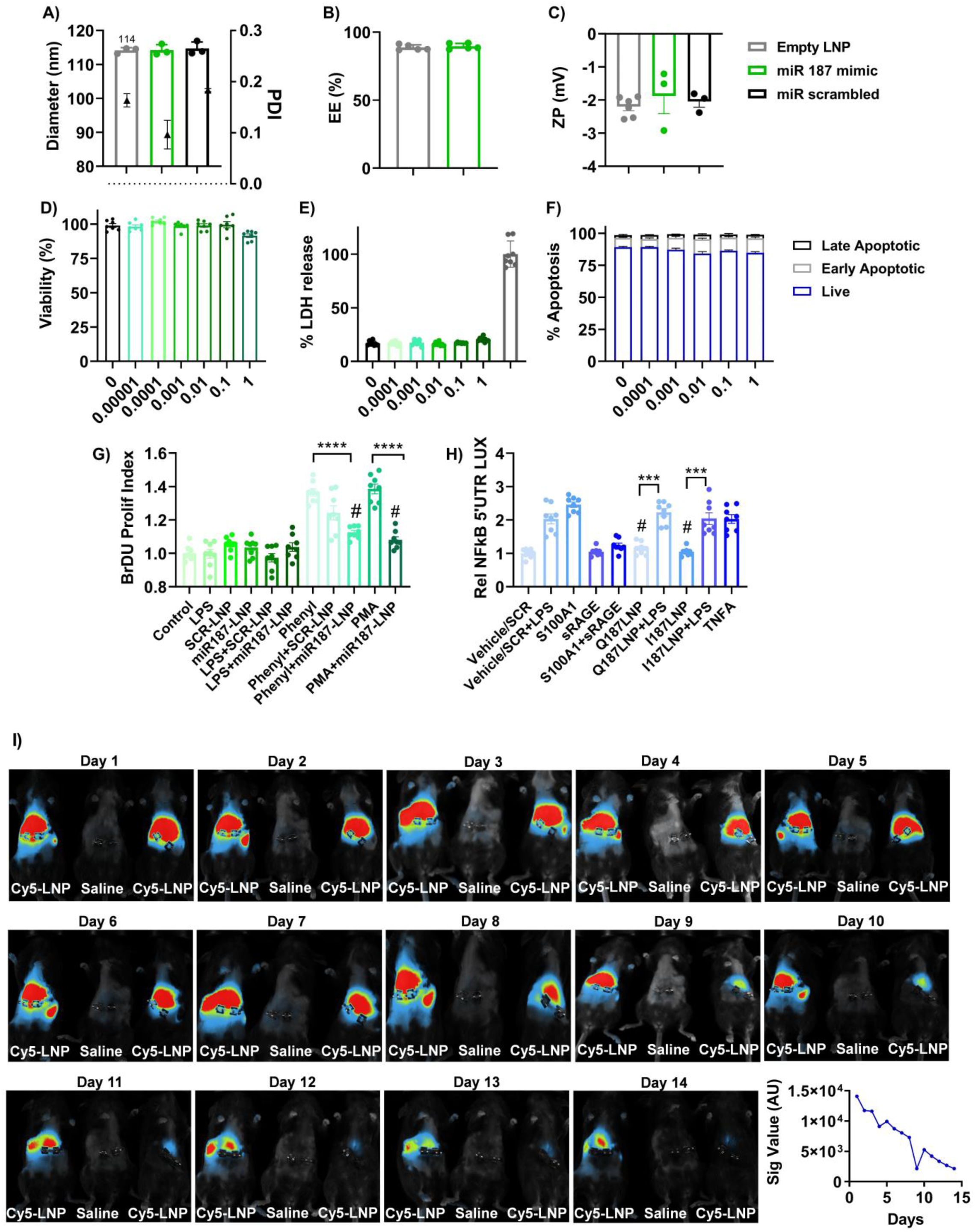
miR-187 LNP characterization, cytotoxicity and persistence of Cy5-labeled lipid nanoparticles (LNPs) in septic mice over 14 days post-injection. Size (A) and polydispersity index (PDI) of LNPs as measured by dynamic light scattering and shown as mm, (B) % encapsulation efficiency (EE) of mimic cargo, and zeta potential (ZP) as mV (C). Cell viability assessed by the Alamar Blue (D), % LDH release (E), and % apoptosis by propidium iodide (PI) staining (F) across a dose range (nmol). Data are presented as mean±SEM. n=3-4/dose. Representative longitudinal in vivo whole-body fluorescence image of Cy5-LNP distribution from Day 1 through Day 14 post-injection (G) and quantification of signal as area under the curve (AU) by densitometry. n=1-2/group.

**Supplemental Figure 5.**
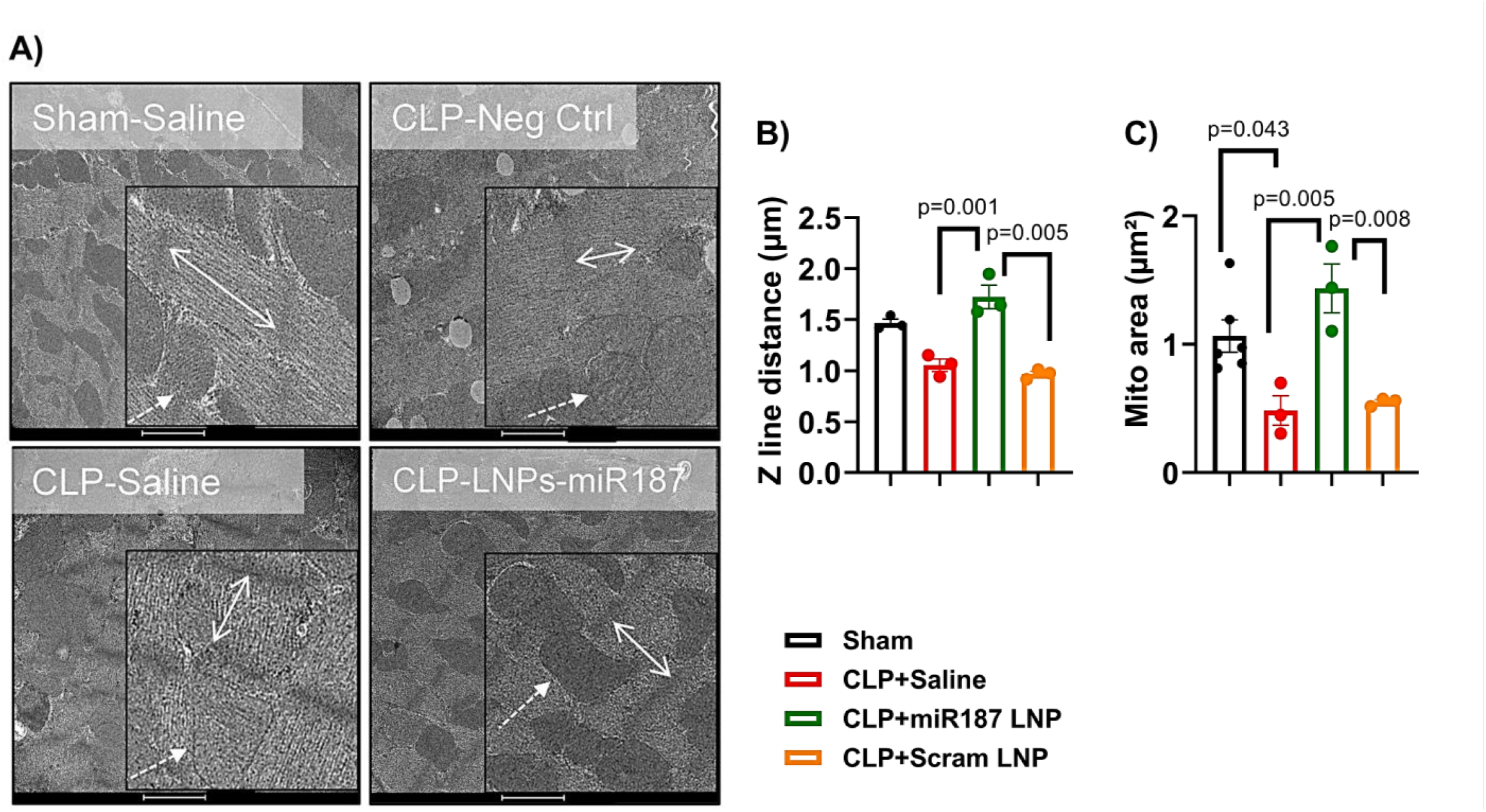
miR-187 LNPs restore sarcomere structure, and mitochondrial integrity in septic mice. Representative transmission electron microscopy (TEM) images of myocardial ultrastructure showing Z-line spacing (arrows) and mitochondrial morphology across groups (B). Quantification of Z-line spacing (C) and mitochondrial area (D). Data are presented as mean ± SD. One-way ANOVA with Bonferroni-Dunn post hoc test was used for statistical comparisons. n=3-4/group.

**Supplemental Figure 6.**
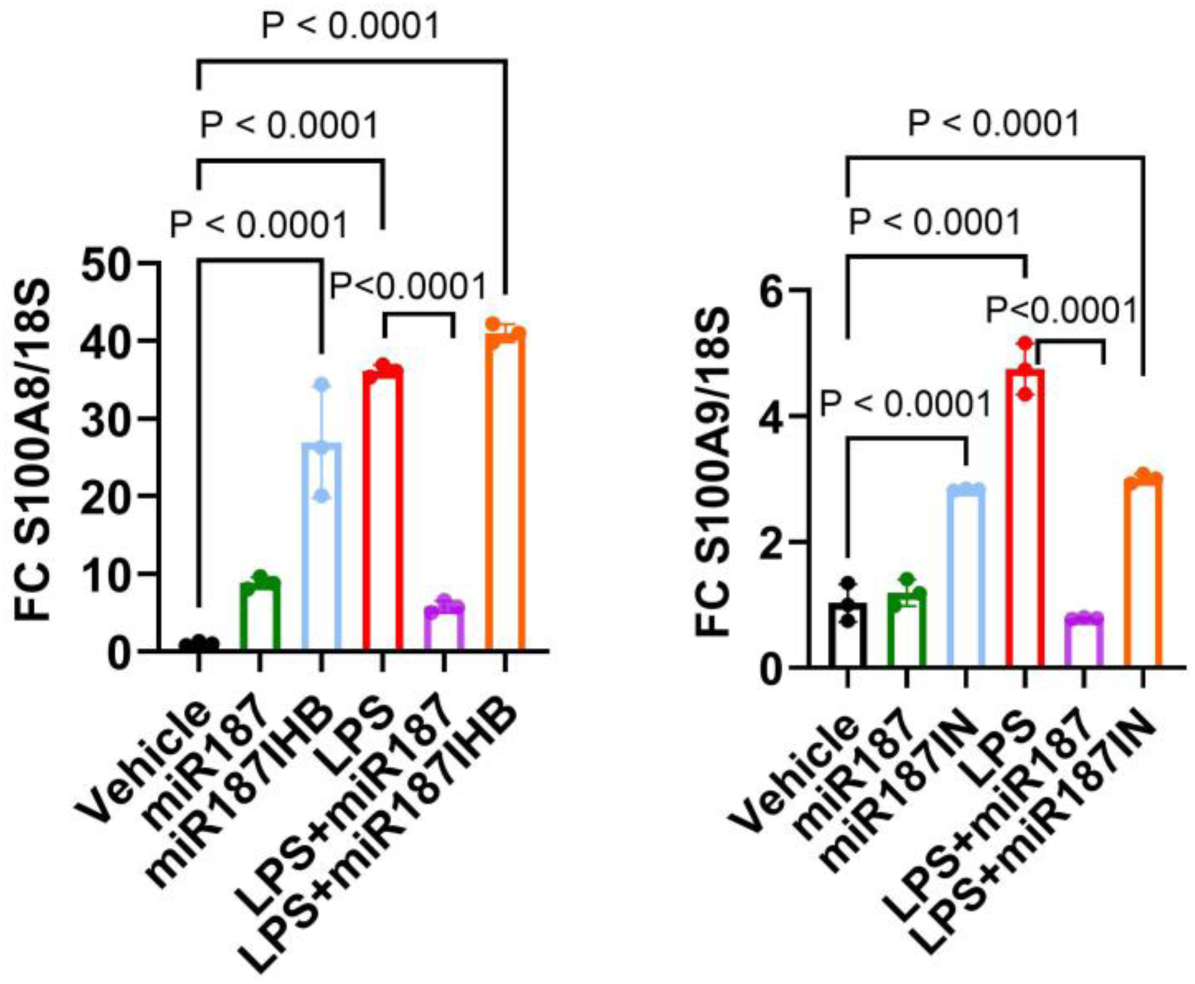
LNP-miR-187 reduces expression of cardiac alarmin genes in adult human cardiomyocytes. AC 10 cells were treated with LNP-miR-187 or LNP-miR-187 inhibitor and 24 hrs later treated with LPS (10 ng/mL) for 24 hrs. Fold change in gene expression of A) S100A8 and B) S100A9 normalized to 18S. Data are presented as mean±SEM. One-way ANOVA followed by Bonferroni-Dunn post hoc test was used. n=3 per group.

**Supplemental Figure 7:**
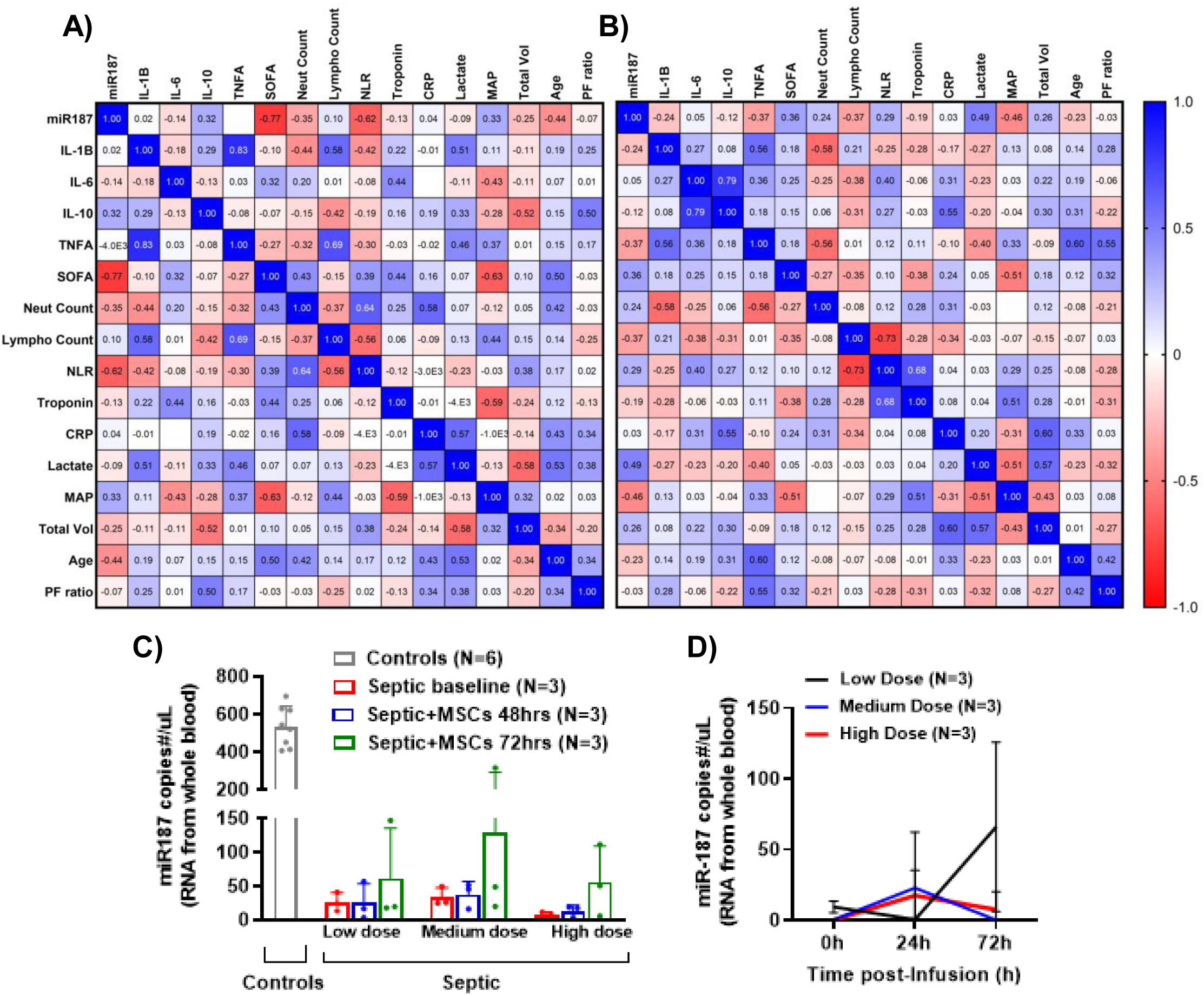
Correlation between miR-187 and clinical features of sepsis and in relation to MSC administration. Pearsons’s correlation matrix of patient characteristics and plasma miR-187 copy number in septic patients with echocardiogram-determined normal (A, n=16) vs. abnormal (B, n=12) cardiac function. Plasma miR-187-3p copy number/μL in septic shock participants that received 0.3 (low dose), 1.0 (mid dose), or 3.0 (high dose) million freshly cultured MSCs per kg body weight, to a maximum of 300 million cells after 24 and 72hrs post-infusion as detected by ddPCR and shown as mean±SD (C-D). One-way ANOVA with Bonferroni-Dunn post hoc test (C-D) were used for statistical comparisons. n=3/group/time point.

## Supplemental Tables

**Supplemental Table 1:** Effect of ciMSC-EVs on blood chemistry analysis of end-organ damage.

| Analyte | Sham |  | CLP Saline |  | CLP Platelets-EV |  | CLP ciMSCs-EV |  |
| --- | --- | --- | --- | --- | --- | --- | --- | --- |
|  | Median | IQR | Median | IQR | Median | IQR | Median | IQR |
| Enzymes |  |  |  |  |  |  |  |  |
| ALT (IU/L) | 28.4 | 14.3 | 41 | 20 | 68.7 | 28.85 | 34.25 | 21.4 |
| AST (IU/L) | 55.4 | 72.6 | 163.7 | 82.9 | 224 | 152.35 | 102.9 | 15.925 |
| ALP (IU/L) | 30.8 | 10.3 | 37.7 | 25.1 | 35.8 | 11.55 | 29 | 6.15 |
| Serum Proteins |  |  |  |  |  |  |  |  |
| Protein, Total (g/L) | 56.6 | 1.8 | 48 | 2.8 | 56 | 6.05 | 53.35 | 3.675 |
| Albumin (g/L) | 33.6 | 1.2 | 23.3 | 1.6 | 27.7 | 3.65 | 27.8 | 1.425 |
| Metabolism |  |  |  |  |  |  |  |  |
| Bilirubin, Total (mg/dL) | 0.15 | 0.11 | 0.29 | 0.01 | 0.24 | 0.08 | 0.2 | 0.1075 |
| Cholesterol, Total (mg/dL) | 78.9 | 60 | 90.4 | 17.4 | 113.8 | 15.7 | 101.5 | 21.725 |
| HDL Cholesterol (mg/dL) | 43.9 | 29.9 | 30.5 | 3.7 | 36.2 | 3.4 | 34 | 3.125 |
| LDL Cholesterol (mg/dL) | 6.59 | 2.94 | 14.09 | 5.35 | 17.17 | 5.99 | 14.235 | 3.11 |
| Triglycerides (mg/dL) | 60.6 | 44.9 | 99 | 32.9 | 98 | 16 | 106 | 9.85 |
| Glucose (mg/dL) | 116.6 | 21.4 | 101.7 | 48.6 | 136.3 | 40 | 124.25 | 37.4 |
| Kidney Function |  |  |  |  |  |  |  |  |
| Creatinine (mg/dL) | 0.206 | 0.002 | 0.156 | 0.009 | 0.206 | 0.054 | 0.1815 | 0.02825 |
| Blood Urea Nitrogen (mg/dL) | 58.5 | 101.9 | 18.7 | 0.8 | 94.3 | 83.2 | 25.35 | 45.4 |
| Serum Electrolytes |  |  |  |  |  |  |  |  |
| Calcium (mg/dL) | 10.678 | 3.169 | 10.038 | 0.063 | 10.902 | 0.44 | 10.522 | 0.5955 |
| Chloride (mmol/L) | 110.4 | 1.8 | 112.8 | 3.5 | 112.6 | 4.45 | 111.2 | 2.675 |
| Phosphorus (mg/dL) | 7.26 | 0.4 | 7.64 | 1.36 | 8.72 | 1.13 | 7.485 | 1.345 |
| Potassium (mmol/L) | 4.58 | 0.75 | 5.41 | 0.29 | 6.09 | 1 | 5.005 | 1.0325 |
| Sodium (mmol/L) | 148 | 3.6 | 148.6 | 0.7 | 148.3 | 3.7 | 150.85 | 0.625 |

**Supplemental Table 2:** Effect of LNP-miR-187 on blood chemistry analysis of end-organ damage.

| Analyte | Sham |  | CLP Saline |  | CLP Neg Ctrl |  | CLP 187 LNP |  |
| --- | --- | --- | --- | --- | --- | --- | --- | --- |
|  | Median | IQR | Median | IQR | Median | IQR | Median | IQR |
| Enzymes |  |  |  |  |  |  |  |  |
| ALT (IU/L) | 45 | 28.7925 | 51.15 | 4.9 | 44.51 | 23.7125 | 62.025 | 30.3475 |
| AST (IU/L) | 85.4 | 13.215 | 219.45 | 59.59 | 159.05 | 89.1 | 195 | 125.695 |
| ALP (IU/L) | 40.38 | 26.98 | 46.7 | 111.5 | 25.45 | 3.645 | 24.83 | 15.175 |
| Serum Proteins |  |  |  |  |  |  |  |  |
| Protein, Total (g/L) | 62.31 | 4.0575 | 54.69 | 10.73 | 50.17 | 7.935 | 52.535 | 4.3175 |
| Albumin (g/L) | 33.985 | 2.995 | 26 | 1.06 | 24.1 | 4.6725 | 24.15 | 5.5025 |
| Metabolism |  |  |  |  |  |  |  |  |
| Bilirubin, Total (mg/dL) | 0.12 | 0.0425 | 0.17 | 0.08 | 0.13 | 0.0675 | 0.115 | 0.0875 |
| Cholesterol, Total (mg/dL) | 139.05 | 16.74 | 103.87 | 5.95 | 104.38 | 23.7725 | 110.35 | 81.48 |
| HDL Cholesterol (mg/dL) | 70.915 | 11.4975 | 36.1 | 11.7 | 40.15 | 10.4575 | 31.9 | 20.93 |
| LDL Cholesterol (mg/dL) | 4.085 | 5.635 | 10.75 | 7.63 | 13.675 | 5.97 | 2.36 | 12.3375 |
| Triglycerides (mg/dL) | 92.06 | 19.7925 | 112.14 | 38.4 | 67.775 | 41.9875 | 88.745 | 70.64 |
| Glucose (mg/dL) | 86.595 | 96.4075 | 66.84 | 56.6 | 62.1 | 71.3625 | 36.365 | 92.565 |
| Kidney Function |  |  |  |  |  |  |  |  |
| Creatinine (mg/dL) | 0.247 | 0.06975 | 0.17 | 0.021 | 0.17 | 0.03325 | 0.176 | 0.07725 |
| Blood Urea Nitrogen (mg/dL) | 28.935 | 1.5025 | 17.38 | 6.2 | 18.4 | 4.405 | 18.12 | 13.8525 |
| Serum Electrolytes |  |  |  |  |  |  |  |  |
| Calcium (mg/dL) | 11.014 | 0.43275 | 9.37 | 1.025 | 10.252 | 0.455 | 9.5565 | 6.8335 |
| Chloride (mmol/L) | 100.86 | 7.9975 | 107.4 | 5.55 | 115.1 | 14.055 | 106.1 | 25.415 |
| Phosphorus (mg/dL) | 6.35 | 2.8275 | 8.72 | 2.19 | 7.47 | 2.2175 | 7.21 | 4.5525 |
| Potassium (mmol/L) | 4.73 | 0.105 | 5.14 | 0.99 | 5.305 | 1.2575 | 4.77 | 2.16 |
| Sodium (mmol/L) | 139.435 | 6.37 | 134.8 | 11.2 | 151.9 | 10.005 | 149.2 | 67.39 |

**Supplemental Table 3:** Pharmacore safety profile of escalating doses of LNP-miR-187.

| Analyte | Control |  | 0.1 nmol miR187 |  | 1 nmol miR187 |  | 3 nmol miR187 |  |
| --- | --- | --- | --- | --- | --- | --- | --- | --- |
|  | Median | IQR | Median | IQR | Median | IQR | Median | IQR |
| Enzymes |  |  |  |  |  |  |  |  |
| ALT (IU/L) | 13.2 | 2.95 | 16.15 | 12.075 | 17.7 | 3.525 | 14.65 | 4.575 |
| AST (IU/L) | 59.1 | 34.05 | 58.1 | 44.8 | 74.05 | 21.875 | 73.15 | 29.65 |
| ALP (IU/L) | 89.5 | 3.75 | 87.1 | 5.425 | 91.95 | 11.65 | 90.8 | 9.4 |
| Serum Proteins |  |  |  |  |  |  |  |  |
| Protein, Total (g/L) | 50.8 | 2.75 | 45.25 | 3.35 | 47.8 | 1.25 | 47.45 | 1.525 |
| Albumin (g/L) | 31.3 | 1.5 | 28.25 | 4.025 | 29.25 | 1.2 | 28.4 | 3.475 |
| Metabolism |  |  |  |  |  |  |  |  |
| Bilirubin, Total (mg/dL) | 0.19 | 0.02 | 0.195 | 0.045 | 0.175 | 0.0325 | 0.17 | 0.015 |
| Cholesterol, Total (mg/dL) | 103.5 | 2.7 | 82.95 | 26.75 | 95.95 | 13.575 | 81.4 | 5.025 |
| HDL Cholesterol (mg/dL) | 58.2 | 2.95 | 45.05 | 17.45 | 48.45 | 3.025 | 44.3 | 1.25 |
| LDL Cholesterol (mg/dL) | 7.36 | 0.71 | 5.16 | 0.32 | 6.835 | 0.83 | 5.465 | 0.93 |
| Triglycerides (mg/dL) | 102.8 | 52.75 | 48.45 | 28.35 | 58 | 16.975 | 84.2 | 38.425 |
| Glucose (mg/dL) | 242 | 20.8 | 255.8 | 54.025 | 297.1 | 45.475 | 266.15 | 41.925 |
| Kidney Function |  |  |  |  |  |  |  |  |
| Creatinine (mg/dL) | 0.17 | 0.041 | 0.23 | 0.044 | 0.222 | 0.05275 | 0.178 | 0.084 |
| Blood Urea Nitrogen (mg/dL) | 27.1 | 3.85 | 24.15 | 4.625 | 21 | 4.65 | 24 | 3.575 |
| Serum Electrolytes |  |  |  |  |  |  |  |  |
| Calcium (mg/dL) | 10.058 | 0.4385 | 9.3595 | 0.6605 | 9.3085 | 0.57225 | 9.365 | 0.56625 |
| Chloride (mmol/L) | 114.2 | 2.15 | 106.6 | 4.325 | 106.4 | 2.45 | 109 | 1.2 |
| Phosphorus (mg/dL) | 8.06 | 1.32 | 7.545 | 1.74 | 8.08 | 2.73 | 7.02 | 1.935 |
| Potassium (mmol/L) | 4.73 | 0.22 | 3.805 | 0.4125 | 3.85 | 0.8375 | 3.88 | 0.2675 |
| Sodium (mmol/L) | 152.1 | 1.25 | 145.6 | 6.05 | 143.35 | 2.275 | 144.7 | 2.2 |
| Blood Cell Counts |  |  |  |  |  |  |  |  |
| WBCs (10<sup>9</sup>/L) | 3.315 | 0.37 | 9.86 | 2.51 | 7.8675 | 0.8575 | 9.6225 | 1.605 |
| Lymphocytes (10<sup>9</sup>/L) | 2.875 | 0.5225 | 8.855 | 2.35 | 6.8925 | 0.87875 | 8.3275 | 1.1375 |
| Neutrophils (10<sup>9</sup>/L) | 0.23 | 0.1175 | 0.775 | 0.135 | 0.675 | 0.11 | 0.7875 | 0.22375 |
| Monocytes (10<sup>9</sup>/L) | 0.065 | 0.045 | 0.13 | 0.02 | 0.125 | 0.05 | 0.1325 | 0.07 |
| Eosinophils (10<sup>9</sup>/L) | 0.08 | 0.0375 | 0.065 | 0.01 | 0.0925 | 0.03125 | 0.0875 | 0.07875 |
| Basophils (10<sup>9</sup>/L) | 0.015 | 0.0275 | 0.03 | 0.01 | 0.045 | 0.0375 | 0.075 | 0.1025 |
| Platelets (10<sup>9</sup>/L) | 926.5 | 804 | 881.5 | 232.25 | 863 | 204.25 | 864 | 150 |

---

**Supplemental Table 4:** Source file showing IntaRNA prediction for binding of miR-187 to regulatory sequences in human and murine MyoD, Tbx15 and Mef2C mRNAs.

